# Structural basis of Cullin-2 RING E3 ligase regulation by the COP9 signalosome

**DOI:** 10.1101/483024

**Authors:** Sarah V. Faull, Andy. M. C. Lau, Chloe Martens, Zainab Ahdash, Hugo Yebenes, Carla Schmidt, Fabienne Beuron, Nora B. Cronin, Edward P. Morris, Argyris Politis

## Abstract

Cullin-Ring E3 Ligases (CRLs) regulate a multitude of cellular pathways through specific substrate receptors. The COP9 signalosome (CSN) deactivates CRLs by removing NEDD8 (N8) from activated Cullins. The structure of stable CSN-CRL can be used to understand this mechanism of regulation. Here we present the first structures of the neddylated and deneddylated CSN-CRL2 complexes by combining single particle cryo-electron microscopy (cryo-EM) with chemical cross-linking mass spectrometry (MS). These structures reveal a conserved mechanism of CSN activation, consisting of conformational clamping of the CRL2 substrate by CSN2/CSN4, release of the catalytic CSN5/CSN6 heterodimer and finally activation of the CSN5 deneddylation machinery. Using hydrogen deuterium exchange-MS we show that CRL2 binding and conformational activation of CSN5/CSN6 occur in a neddylation-independent manner. The presence of NEDD8 is required to activate the CSN5 active site. Overall, by synergising cryo-EM with MS, we identified novel sensory regions of the CSN that mediate its stepwise activation mechanism and provide a framework for better understanding the regulatory mechanism of other Cullin family members.

**One sentence summary:** Structure and dynamics of the CSN-CRL2 complexes assessed by cryo-electron microscopy and structural mass spectrometry.

## MAIN

Cullin-RING Ligases (CRLs) are modular, multi-subunit complexes that constitute a major class of ubiquitin E3 ligases^1,2^. CRLs coordinate the ubiquitination of substrates as either a signal for degradation via the 26S proteasome, or to alter the function of the target protein^2,3^. The CRL2 E3 ligase consists of a Cullin-2 (CUL2) scaffold in association with a catalytic RING-box protein (RBX1), with the substrate adaptors Elongin B (ELOB) and C (ELOC) at its N-terminal^4^. When associated with the von Hippel-Lindau (VHL) tumour suppressor substrate receptor, the CRL2 complex is the primary regulator of the Hypoxia Inducible Factor 1-α (HIF-1α) transcription factor^5,6^. Mutations in the interface between VHL, ELOB and ELOC can deactivate CRL2 leading to an accumulation of HIF-1α, which can in turn drive tumorigenesis through the over-activation of oncogenes^7^. Moreover, CRL2 has recently been identified as a potential target for small molecular inhibitors and PROTACs – a new class of cancer drugs that promote degradation of tumorigenic gene products^8^-^10^. These fascinating systems have been described in detail by a number of excellent reviews^1,2,3^.

Activation of CRL2, in common with other members of the CRL family, involves a cascade of E1, E2 and E3 enzymes, which conjugate the ubiquitin-like protein NEDD8 (N8) to residue K689 on the Cullin-2 scaffold^11^. In its activated state, CRL2∼N8 (the ∼ stylization denotes a covalent interaction) recruits the ubiquitin-conjugated E2 enzyme via the RING domain of RBX1^12^. Ubiquitination now takes place, covalently adding ubiquitin to the substrate molecule docked at the CRL2 N-terminal. The activity of CRL2 is negatively regulated by the 331 kDa Constitutive Photomorphogenesis 9 Signalosome (CSN) complex, frequently referred to as the COP9 signalosome complex^13^-^15^. The CSN was originally identified as consisting of eight subunits (designated as CSN1-8 by decreasing molecular weights of 57-22 kDa), and is organized in a “splayed hand” architecture, which has high sequence and structural homology to the proteasome lid^13,14,16^-^18^. CSN1, 2, 3, 4, 7 and 8 are structurally homologous to each other and together contribute to the fingers of the “splayed hand” which arise from extended N-terminal α-helical repeats^16,18^. CSN5 and 6 are also closely related structurally and form a globular heterodimer located on the palm of the hand. CSN5 is responsible for the deneddylase activity of the CSN. A ninth subunit, CSNAP, has recently been identified, and is thought to play a role in stabilising the CSN complex^18^.

Electron microscopy (EM) based structural analysis has provided important insights into the mechanism of CRL1 regulation by CSN^16,19^. CRL4A and CRL3 have also been observed to form such complexes^20^. However, despite intense interest, structural information of the CSN bound to CRLs remains limited to CRL1^12,16,19^, CRL4A^20^, and a low-resolution map of a dimeric CSN-CRL3-N8^20^complex. In particular the analysis of the CSN-CRL4A complex^20^identified at least three major steps by which CRL∼N8 is deneddylated by the CSN. In the first step, the extended N-terminal helical modules of CSN2 and CSN4 conformationally “clamp” the C-terminal domain of the CRL4A∼N8 and RBX1^16,19,20^. The second step involves the release and consequent relocation of CSN5/CSN6 closer to NEDD8, brought about by disruption of the CSN4/CSN6 interface^20^. Disrupting the binding interface between CSN4/CSN6 through removal of the CSN6 insertion-2 loop (Ins-2), resulted in enhanced deneddylase activity^13^, presumably due to more complete release of CSN5/CSN6. In the final step, the mobile CSN5 binds to NEDD8, leading to deneddylation via its JAB1/MPN/MOV34 (JAMM) metalloprotease domain^21^. The JAMM motif consists of H138, H140 and D151 zinc-coordinating residues, and residue E104 of the CSN5 insertion-1 loop (Ins-1)^13^. In apo-CSN, the Ins-1 loop occludes the CSN5 active site, auto-inhibiting the deneddylase^13,22^. Deneddylation is also severely diminished by a H138A point mutation in CSN5^13^.

Surprisingly, the CSN can also form complexes with each of the Cullin 1-5 family members even without NEDD8^23^. Free CRLs such as CRL1 have been reported to readily bind and inhibit the CSN, albeit at relatively lower affinity than the neddylated CRL1^24^. While the exact role of CSN-CRL complexes remains unclear, it has been hypothesised that these complexes may function to regulate the cellular level of ubiquitin ligase activity of CRLs once they have been deneddylated, effectively sequestering E3 ligases from the intracellular environment^24^.

Building on the existing knowledge of the CSN-CRL systems, here we pose the question: are similar structural changes to be found in other CSN-CRL complexes, and how does binding of neddylated CRLs lead to activation of the CSN5 catalytic site? To address this, here, we present novel structures of the CSN-CRL2∼N8 complex, together with the first structure of the CSN-CRL2 deneddylation product. We complemented our cryo-EM analysis with chemical cross-linking mass spectrometry (XL-MS) allowing us to clarify the positions of particularly dynamic regions in the complexes. We used hydrogen-deuterium exchange mass spectrometry (HDX-MS) to interrogate the role of the CSN4/CSN6 interface in communicating CRL binding to the CSN5 active site. Overall, our structures of the CSN-CRL2∼N8 and its deneddylation product, the CSN-CRL2, provide a molecular level understanding of how deneddylation is performed by the CSN.

## RESULTS

### Cryo-EM structures of the CSN-CRL2∼N8 complex

To study the molecular interactions between neddylated CRL2 (CRL2∼N8) and the CSN, we performed single-particle cryo-EM to resolve a structure of the assembled CSN-CRL2∼N8 (referred to as the holocomplex) (**Fig. S1-S2**). The mutation H138A in the catalytic site of CSN5 subunit makes it possible to assemble CSN-CRL2∼N8 complexes in which NEDD8 remains covalently attached over the time scale for experimentation^16,19^. This mutant form of CSN was used throughout the work described below, unless otherwise specified.

Using three-dimensional classification, we were able to generate maps of three different structures: a) a holocomplex map at 8.2 Å, b) a map of the complex with little or no density for VHL at 8.0 Å, and c) a map of the complex with little or no density for CSN5/CSN6/VHL at 6.5 Å (**Fig. S2**, **Fig. S3**). The two partial complexes likely arise from compositional heterogeneity in the original samples from which the structural analysis has succeeded in isolating subpopulations.

Next, we fitted into each map, the crystallographic CSN (PDB 4D10) and a homology model of the CRL2 (including the VHL-ELOB-ELOC) using molecular dynamics flexible fitting^25^(**Table 1; Methods**). In the model of the holocomplex (8.2 Å), the main interactions occur between the C-terminal end of Cullin-2 and the extended N-terminal helical repeats of CSN2 and CSN4 (**Fig. 1a-b**). Compared to their apo-CSN crystal structure conformations (PDB 4D10), CSN2 and CSN4 are moved by 30 and 51 Å, respectively, towards Cullin-2 (**Fig. S4a**). In each case, the motion is a swinging rotation about hinges located close to the CSN2 and CSN4 winged helix domains: in the case of CSN2 this is coupled with an additional rotation about the axis of the superhelix formed from its N-terminal helical repeats. In the case of CSN4 the movement is coupled to the detachment of CSN4 from the Ins-2 loop of CSN6 by ∼30 Å and leads to a ∼12 Å shift in CSN5 (**Fig. 1c**, **Fig. S4b**). Only minor conformational changes were found in the CSN1, CSN3, CSN7B or CSN8 subunits. In the CRL2∼N8 moiety, a number of relatively small rearrangements of Cullin-2, RBX1, ELOB, ELOC and VHL subunits were observed compared to its crystal structure^26^(**Fig. S4c-d**).

**Table 1.**
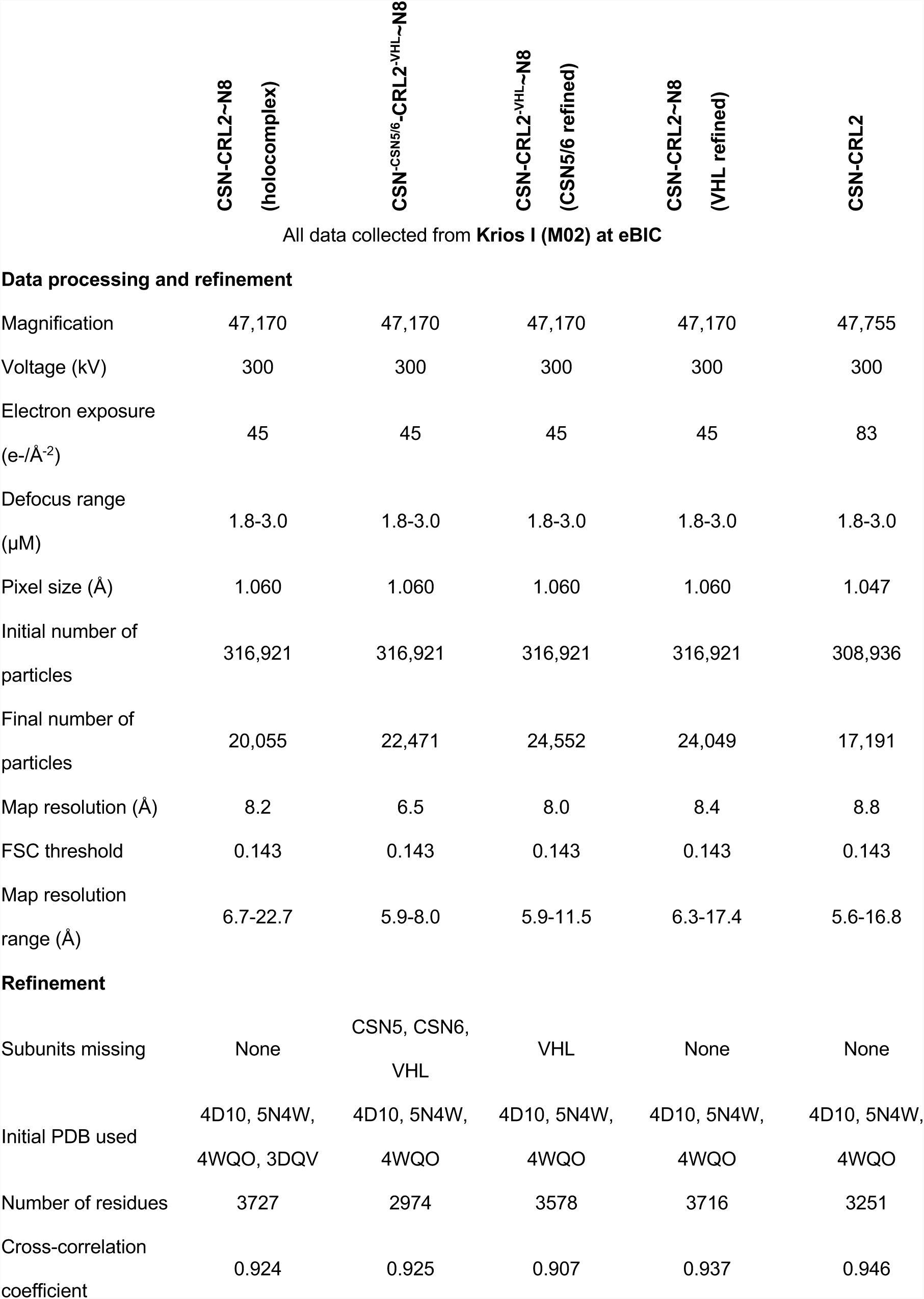
Cryo-EM refinement parameters.

**Fig. 1.**
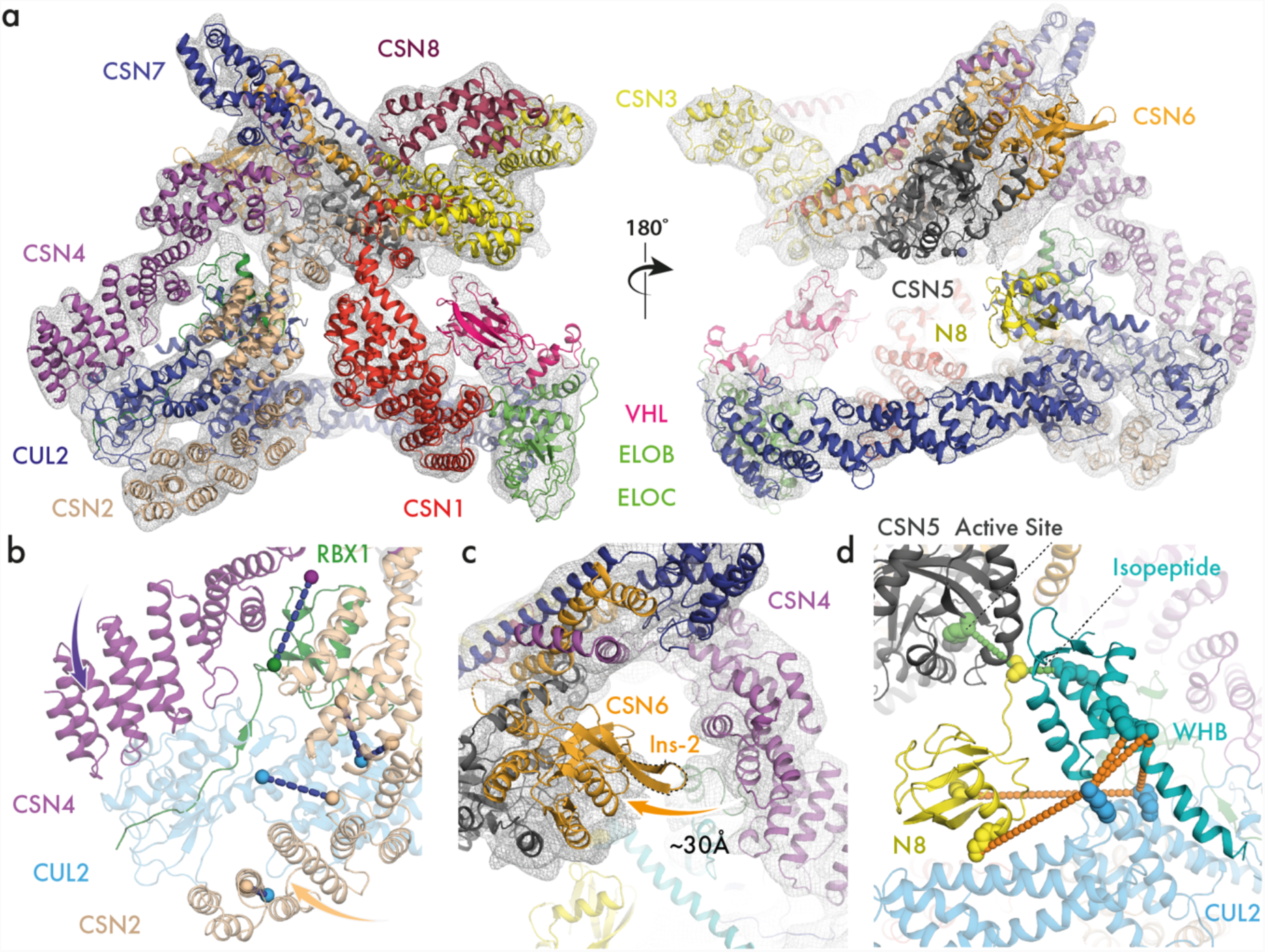
Cryo-electron microscopy structures of the CSN-CRL2∼N8 complex. (**a**) Molecular model of CSN-CRL2∼N8 fitted into cryo-EM density (8.2 Å resolution) from front and back views. (**b**) Conformational clamping of CRL2 by CSN2 and CSN4. Cross-links shown are between CSN4-RBX1 (CSN4^K200^-RBX1^K105^, purple-green spheres), and four between CSN2-CUL2 (CSN2^K157^-CUL2^K489^, CSN2^K263^-CUL2^K462^, CSN2^K225^-CUL2^K462^, CSN2^K64^-CUL2^K404^, beige-blue spheres). (**c**) View showing ∼30Å separation of CSN6 Ins-2 loop from CSN4 following CRL2∼N8 binding. (**d**) Modelled position of WHB∼N8 using cross-links of the CSN-CRL2∼N8 (see also **Fig. S5d**).

In our EM maps and the other published CSN-CRL structures^16,19,20^, the exact position of NEDD8 and the Cullin-2 Winged-Helix B (WHB) domain were difficult to determine. To address this limitation, we carried out XL-MS experiments on the CSN-CRL2∼N8 complex using the bis(sulfosuccinimidyl)suberate (BS3) cross-linker which targets lysine sidechains (**Methods**). We identified a total of 24 inter-and 60 intra-protein cross-links (**Table S1, Fig. S5a-b**). To generate a model of the CSN-CRL2∼N8, we performed cross-link guided modelling which allows the placement of the WHB, NEDD8 and VHL subunits using identified cross-links from XL-MS (**Methods**). We imposed a cross-link distance threshold of 35 Å which takes into account the length of two lysine side chains (15 Å), the BS3 cross-linker length (10 Å) and an extra 10 Å to allow for domain-level flexibility (**Methods**). Our model of the CSN-CRL2∼N8 satisfies all cross-link distances (**Fig. S5c**). Three cross-links between Cullin-2-WHB (Cullin-2^K382^-WHB^K720^, Cullin-2^K382^-WHB^K677^and Cullin-2^K433^-WHB^K677^) were used for the positioning of the WHB domain (**Fig. S5d**, red text). A further two cross-links between Cullin-2 and NEDD8 (Cullin-2^K382^-N8^K33^and Cullin-2^K433^-N8^K6^) allowed us to place NEDD8 near CSN5 (**Fig. 1d, Fig. S5d**, green text). In this conformation, the isopeptide bond of NEDD8 is juxtaposed towards to the CSN5 active site. For the isopeptide bond of NEDD8 to reach the CSN5 active site, the Cullin-2 WHB domain must be extended from its crystallographic conformation towards the CSN5 by 19 Å (**Fig. S5e**).

### Structure of the deneddylated CSN-CRL2 complex

Having determined the structure of the CSN-CRL2∼N8 complex, we next sought to detail any conformational differences in the deneddylated CSN-CRL2. The affinity of CSN for the non-neddylated CRLs is significantly lower than for the neddylated forms, limiting the yield of the desired product^19^. We used native MS experiments to confirm the formation of the complex (**Fig. S6**). We next resolved a cryo-EM map of the CSN-CRL2 to 8.8 Å resolution **(Fig. S7**). Although the resolution of the CSN-CRL2 map is similar to that for CSN-CRL2∼N8 holocomplex, we only observed partial density for VHL and CSN4. This is most likely due to the reduced stability of the CSN-CRL2 assembly in the absence of NEDD8. We were however able to use the density obtained to model the CSN-CRL2 structure, which has principally the same topology as the neddylated holocomplex (**Fig. 2a**). Using the same procedure as for the neddylated CSN-CRL2∼N8 complex, we fitted CSN and CRL2 subunits into the density map of the CSN-CRL2 complex (**Methods**). We then utilised cross-link guided modelling to establish the position of the WHB which lacked clear density, similar to the neddylated holocomplex (**Fig. S8; Methods**).

**Fig. 2.**
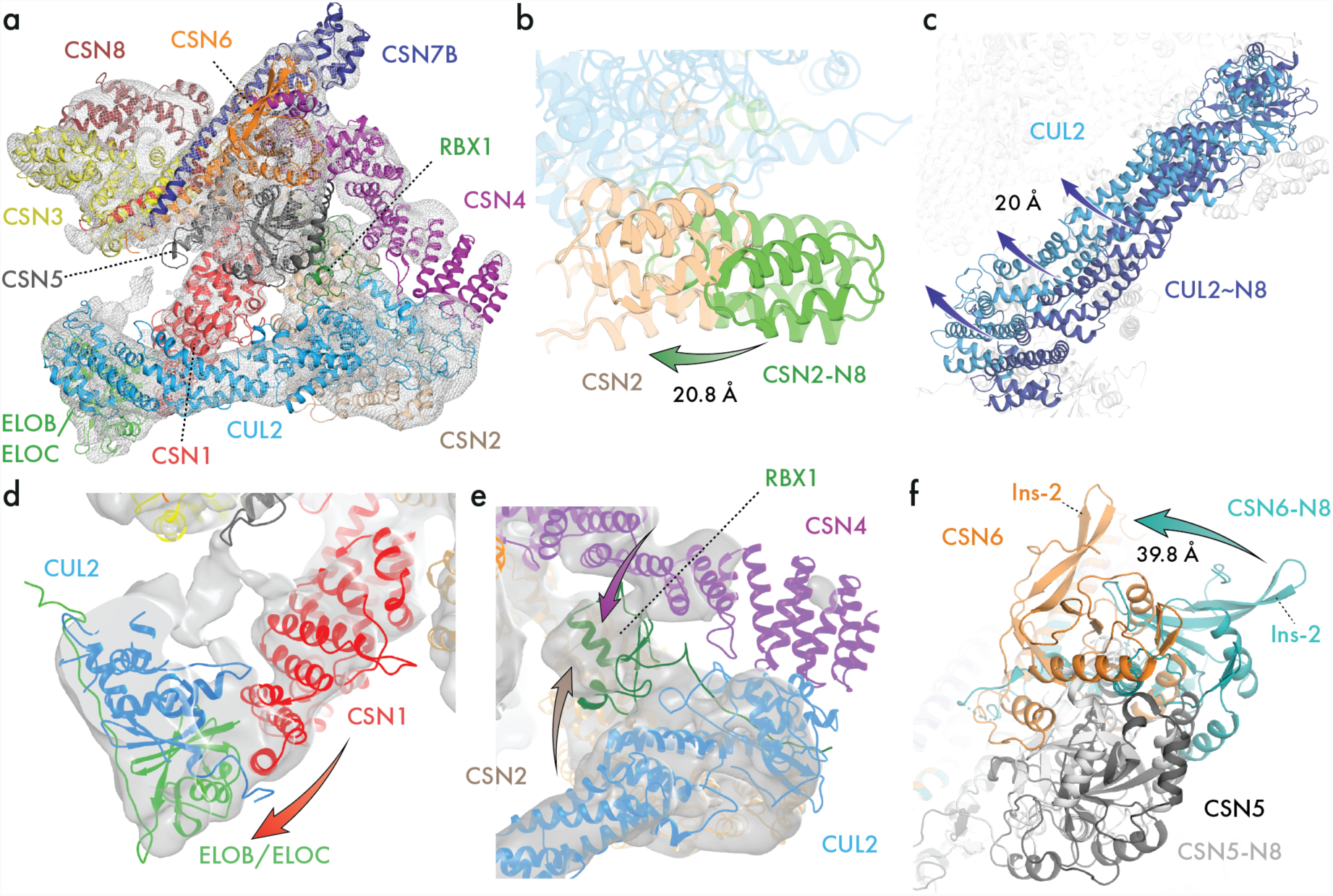
Structure of the deneddylated CSN-CRL2 complex. (**a**) The fitted density map of deneddylated CSN-CRL2 structure determined by combining cryo-EM and XL-MS. Alignment of the CSN C-terminal helix bundle from the neddylated and deneddylated holocomplexes reveals differences in (**b**) CSN2 and (**c**) Cullin-2. (**d**) CSN1-ELOB interface established from the rotation of Cullin-2 in (**c**). (**e**) RBX1 is clamped between CSN2 and CSN4. (**f**) Conformational changes in CSN6 in the absence of NEDD8.

To determine any local changes across the CSN-CRL2∼N8 in the absence of NEDD8, we aligned the cryo-EM models of neddylated and deneddylated holocomplexes using the C-terminal helical bundle as a reference point (**Fig. S9**). We systematically compared the conformations of each subunit (**Fig. S9b-i**). Compared to its structure in the CSN-CRL2∼N8, the N-terminal helices of CSN2 are shifted by ∼21 Å towards the Cullin-2 C-terminal domain (**Fig. 2b, Fig. S9c**). This change in CSN2 in the absence of NEDD8, leads to a structural difference in Cullin-2 which rotates upwards towards the rest of the CSN by 20 Å (**Fig. 2c, Fig. S9a**). The position adopted by Cullin-2 in the deneddylated holocomplex, places ELOB closer to CSN1, forming a CSN1-ELOB interface (**Fig. 2d**). Interactions between substrate adaptor complex and CSN1 have similarly been reported for the CSN-CRL1∼N8^16^and CSN-CRL4A∼N8^20^. RBX1 remains clamped between CSN2 and CSN4 (**Fig. 2e**).

The most striking conformational differences were observed in CSN6 (**Fig. 2f, Fig. S9g**). In the absence of NEDD8, CSN6 is dramatically shifted away from its position in the neddylated holocomplex by ∼40 Å (**Fig. 2f**). This previously unknown conformation of CSN6 appears to be unique, differing from the conformation captured in our neddylated holocomplex, and the CSN-CRL4A∼N8 and CSN-CRL3∼N8^19^structures. Similar to the neddylated holocomplex, no significant changes were identified in CSN3, CSN7B and CSN8 subunits. Overall, comparison between the CSN-CRL2∼N8 and CSN-CRL2 structure reveal significant conformational rearrangements in CSN5/CSN6 and the N-terminal domain of CSN2.

### HDX-MS reveals a stepwise mechanism of CSN activation

Having determined the structures of the neddylated and deneddylated CSN-CRL2 complexes, we set off to characterise the local dynamics using HDX-MS. HDX-MS provides peptide-level information on the dynamics of proteins through monitoring the exchange events of amide hydrogens for bulk deuterium in the surrounding solution environment^26-31,32^. Here, we performed a set of two differential HDX-MS experiments to determine the effect of: a) CRL2∼N8 binding to CSN, denoted as Δ(CSN-CRL2∼N8 – CSN), and b) CRL2 binding to CSN, denoted as Δ(CSN^WT^-CRL2 – CSN^WT^) (**Fig. 3a, Fig. S10-11**). Regions that exhibit significant HDX differences brought about by the addition of the ligand (i.e. CRL2 and CRL2∼N8) are labelled as stabilising (negative ΔHDX; coloured blue) or destabilising (positive ΔHDX; coloured red). It is important to note that our HDX-MS experiments did not cross-compare the CSN^5^H^138A^and CSN^WT^ complexes.

**Fig. 3.**
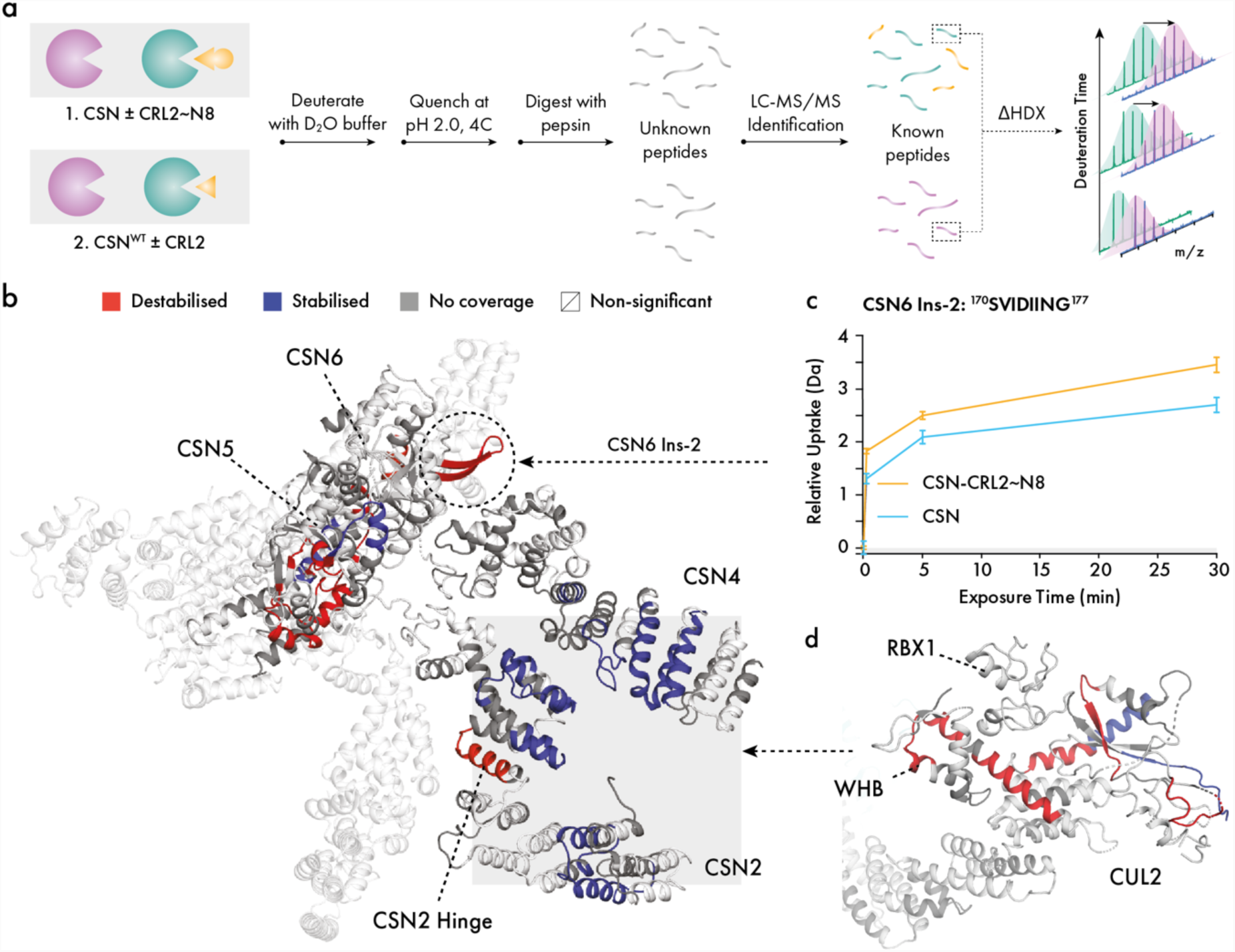
Effect of NEDD8 on the CSN4/CSN6 interface. (**a**) Investigating the effect of CRL2∼N8 and CRL2 binding on CSN and CSN^WT^, respectively. The two experiments involve deuterating the complexes for 0.25, 5 and 30 min timepoints, a quench step to halt the deuteration, and digestion to the peptide level. Peptides are then identified using liquid chromatography-tandem mass spectrometry (LC-MS/MS) and a database search. (**b**) Effect of CRL2∼N8 binding on the CSN. (**c**) Relative deuterium uptake over 30 minutes for the CSN6 Ins-2 peptide (^170^SVIDIING^177^). (**d**) Effect of CSN on the C-terminal domain of CRL2∼N8. Colour scheme follows that of (b).

In both Δ(CSN-CRL2∼N8 – CSN) and Δ(CSN^WT^-CRL2 – CSN^WT^) experiments, extensive regions in the N-terminal helices of CSN2, CSN4 and the globular domain of RBX1 exhibited stabilisation upon the incubation of CSN with its CRL2∼N8 and CRL2 substrates (**Fig. 3b; Fig. S12a-b**). These observations are in line with the conformational clamping by CSN2/CSN4 onto the C-terminal of CRL2 as seen in the cryo-EM structures of neddylated and deneddylated complexes. Within CSN2 of both experiments, we observed significantly destabilised regions around helical modules 6-9. These observations likely identify the hinge points which permit the bending of CSN2 to clamp onto the Cullin-2 C-terminus (**Fig. 3b; Fig. S12a-b**).

### Conformational remodelling of the CSN5 active site is achieved in the presence of NEDD8

We next considered the release mechanism of the CSN5/CSN6 subunits of both the neddylated and deneddylated holocomplexes. In both Δ(CSN-CRL2∼N8 – CSN) and Δ(CSN^WT^-CRL2 – CSN^WT^) experiments, the Ins-2 loop of CSN6 was destabilised, correlating with the release of CSN6 from its interface with CSN4 and in line with the allosteric activation mechanism of CSN by CRL4A^19^(**Fig. 4a-b i**). An interesting difference between the neddylated and deneddylated complexes is that the CSN6 α4 helix is destabilised only in the absence of NEDD8 (**Fig. 4b i**). Similarly, the CSN5 α7 helix is also destabilised in both neddylated and deneddylated conditions (**Fig. 4a-b ii**). The CSN6 α4 and CSN α7 helices are topologically knotted in the CSN5/CSN6 heterodimer and tether the globular domains of CSN5/CSN6 to the C-terminal helical bundle^13^. These observations suggest that structural changes are required in the helical knot to bring about release of the CSN5/CSN6 globular domains from their apo conformation.

**Fig. 4.**
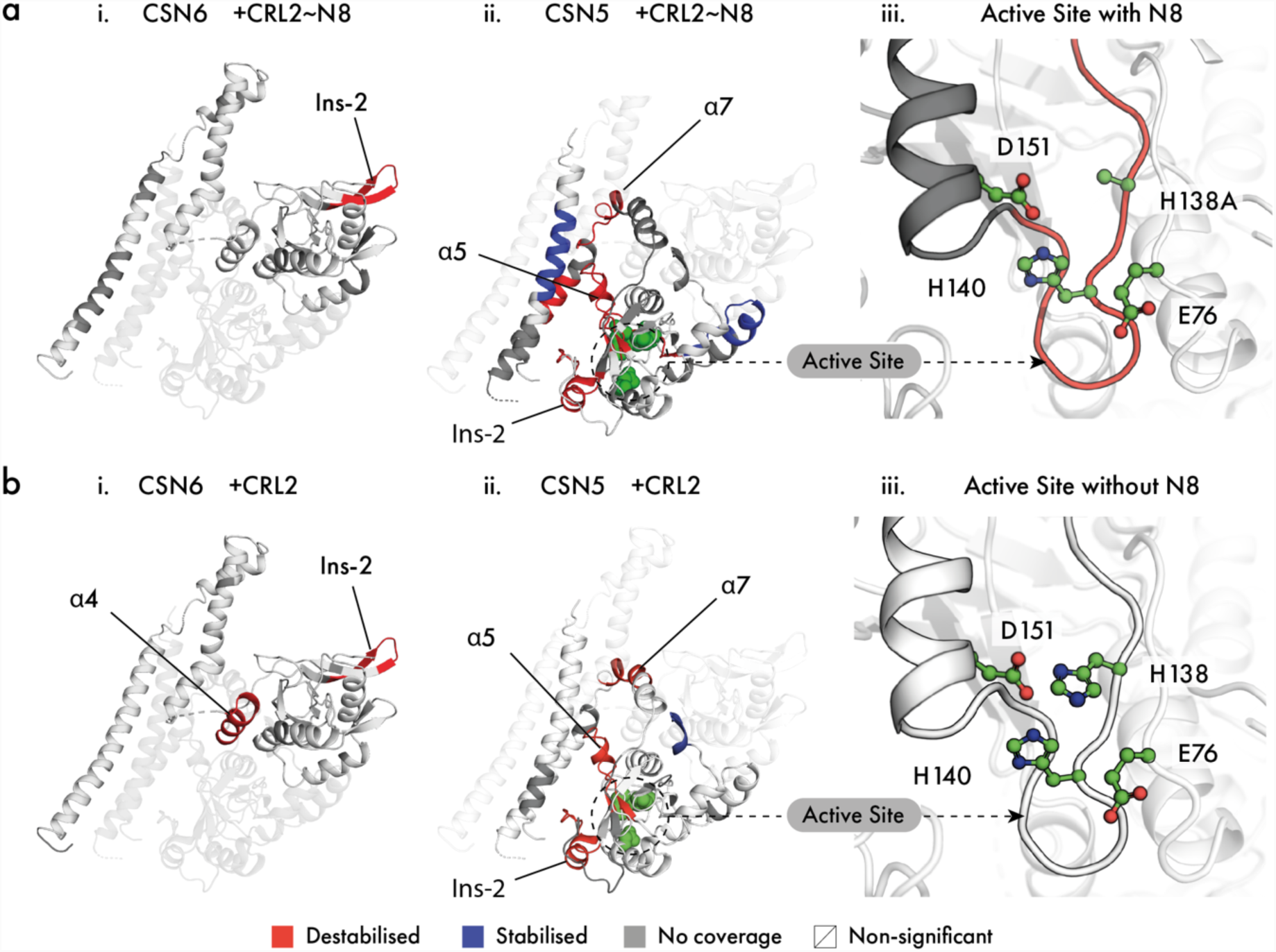
Conformational response of CSN5/CSN6 to NEDD8. Differential HDX-MS of (**a**) Δ(CSN-CRL2∼N8 – CSN) and (**b**) Δ(CSN^WT^-CRL2 – CSN^WT^). Regions exhibiting significant deuterium uptake differences in (**i**) CSN6, (**ii**) CSN5 and (**iii**) CSN5 active site are highlighted in red for destabilised and blue for stabilised areas. Regions without coverage or exhibit non-significant changes are in grey and white respectively.

Another finding is that we identified destabilisation in the Ins-2 loop of CSN5 (**Fig. 4a-b ii**). The Ins-2 loop of CSN5 has a lesser understood role in CSN activation. In isolated CSN5, the Ins-2 loop is highly disordered^22^(**Fig. S13a**), while it folds into a helical-loop structure when incorporated into the CSN^13^(**Fig. S13b**). Accompanying the changes in the CSN5 Ins-2 loop, in both comparative HDX-MS experiments, we detected destabilisation of the α5 helix area which surrounds the CSN5 active site (**Fig. 4a-b ii**). The changes in both the CSN5 Ins-2 and α5 helix indicate a major conformational remodelling in the area adjacent to the CSN5 active site, which can be triggered through the binding of both CRL2 or CRL2∼N8 to the CSN in a NEDD8-independent manner. It is only in the presence of NEDD8, that the CSN5 active site is further destabilised suggesting that in a final activation step, NEDD8 induces conformational changes in the active site itself (**Fig. 4a-b iii, Fig. S14**).

## DISCUSSION

Here we have combined EM and MS analyses to provide new insights into the mediation of CRL2 by the CSN. We have described the first structures of CSN-CRL2∼N8 and a novel deneddylated CSN-CRL2 structure. As far as we are aware, this is also the first structural description of a non-neddylated CSN-CRL complex. Furthermore, we combined cryo-EM maps with comparative HDX-MS to expand on the stepwise activation mechanism of the CSN, involving a conformational network of both NEDD8-independent and dependent stages. We suggest that the steps which lead to deneddylation are mostly NEDD8-independent, except for the remodelling of the CSN5 active site which requires NEDD8 to encounter the CSN5 active site.

Our map of the deneddylated CSN-CRL2 holocomplex represents a complex in which the CSN is still associated with its CRL2 reaction product. Resolving this structure has provided several important details into how activation of the CSN is achieved. Our comparison of the neddylated and deneddylated holocomplex structures indicated that the CSN2 contacts the CRL2 C-terminal domain in a slightly different conformation to when the CRL2 is modified with NEDD8. Between both neddylated and deneddylated conformations, the clamping by CSN2 involves destabilisation of the CSN2 helical modules 6-9 which function possibly as a hinge that allows the CSN2 to bend upwards towards the CRL2. The plasticity of these N-terminal helices presumably permits the binding of deneddylated and alternative Cullin isoforms. HDX-MS further indicates that RBX1 and CSN4 form an interface, which is more prominent in the absence of NEDD8, as shown through stabilisation of the two interfaces (**Fig. S12b**). Overall, the conformational variations seen in the CSN2 N-terminal helical modules (**Fig. 2b**), the bend of CRL2 (**Fig. 2c**) and the HDX differences in CSN4/RBX1 (**Fig. S12b**) may therefore be attributable to the seemingly promiscuous affinity that allows CSN2 to bind to each of the different Cullins regardless of neddylation.

We uncovered structural and dynamical aspects of both the neddylated and deneddylated CSN-CRL2 complexes. Beginning our interpretation from valuable studies of the CSN-CRL1∼N8 and CSN-CRL4A∼N8 systems, we have identified several major conformational switches of the CSN which must be activated by the CRL2 substrate to bring about deneddylation. While some of these steps are conserved ubiquitously among other CSN-CRL complexes (e.g. CSN2/CSN4 clamping in CSN-CRL1∼N8 and CSN-CRL4A∼N8), our study has identified additional steps for the CRL2-bound CSN complex, enabling the most complete mechanistic interpretation of CSN activation by a CRL to date (**Fig. 5**). In the first activation step, the CSN and CRL2∼N8 associate through major conformational changes in CSN2 and CSN4, which clamp onto the CRL2 (**Fig. 5a-b**). The dramatic change in CSN4 breaks its interface with CSN6 through the CSN6 Ins-2 loop and releases the CSN5/CSN6 heterodimer (**Fig. 5c**). The release of CSN5/CSN6 appear consistent with the destabilised knotted helices of CSN6 that our HDX has identified. It is plausible that these two helices function as the mechanical hinges which allow the CSN5/CSN6 to be released from their auto-inhibited conformations but remain tethered to the rest of the CSN. Although the resolution presented by CSN5 in our cryo-EM structures prevents us from making molecular level observations, our HDX data can provide local detail for the CSN5 active site. The release of CSN5/CSN6 is accompanied by HDX changes in areas surrounding the CSN5 active site, including the CSN5 Ins-2 loop. Up to here, the changes experienced by the CSN can be brought about in a NEDD8-independent manner. In the next stage, the presence of NEDD8 acts as a selectivity filter which results in remodelling of the CSN5 active site itself (**Fig. 5d**). These changes presumably expose the CSN5 JAMM ligands of the metalloprotease site and allow subsequent deneddylation to occur (**Fig. 5e**). Finally, deneddylation ensues with the cleavage of NEDD8 from CRL2 and the dissociation of the complex (**Fig. 5f**). The fact that CSN can then re-associate with its CRL2 reaction product following dissociation, as shown by our study and structure of the CSN-CRL2, suggests that this complex may serve a lesser understood role in the CSN-CRL network.

**Fig. 5.**
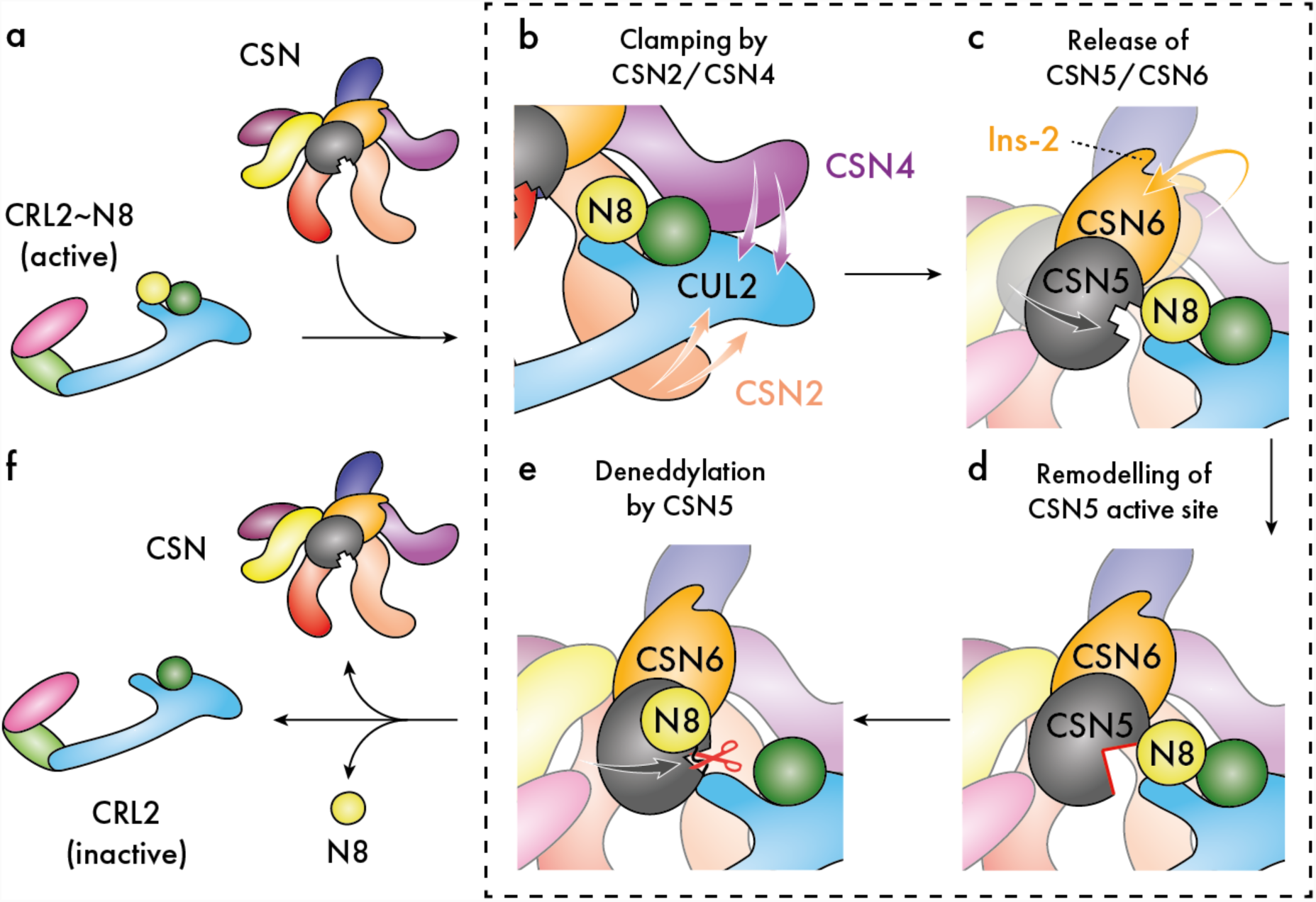
Schematic of CRL2 regulation by the CSN. (**a**) Neddylated (active) CRL2∼N8 is regulated by the CSN. (**b**) CRL2∼N8 is bound by the CSN, principally through clamping by the extended N-terminal helical modules of subunits CSN2 and CSN4. (**c**)The topological changes that occur in CSN4 upon CRL2 binding result in detachment of the CSN6 Ins-2 loop and subsequent release of the CSN5/CSN6 heterodimer in a NEDD8-independent step. (**d**) The presence of NEDD8 triggers remodelling of the CSN5 active site, (**e**) permitting deneddylation of CRL2. (**f**) Cleavage of NEDD8 results in inactive CRL2 and the CSN dissociation.

Furthermore, there may be additional roles for the CSN as suggested by the compositional plasticity seen in our CSN-CRL2∼N8 classes. In our map of the CSN-CRL2∼N8, the Cullin arm was observed to shift downwards away from CSN3 in maps deficient of the VHL substrate receptor (**Fig. S15**), consistent with the coupling of VHL binding to a conformational change in the rest of the CRL2 portion of the protein. This type of behaviour may allow the CRL2 to adapt to individual substrates and substrate receptors which vary in size and geometry as has been suggested with the CRL4A system^20^. Additionally, it may reflect changes associated with remodelling of the CRL2 by the dissociation of the VHL and the binding of alternative substrate receptors. In future work we will seek to determine whether the CSN can mediate substrate receptor dissociation.

Overall, our study has provided unprecedented level of detail at the role of CRL2 and NEDD8 on regulating the activation mechanism of CSN. We have revealed that the series of mechanistic responses of the CSN that lead up to deneddylation, can be triggered even by the CRL2 reaction product in a completely NEDD8-independent manner. The presence of NEDD8 on the activated CRL2 substrate, triggers remodelling within the catalytic site of the CSN5 subunit during the final stage of CSN activation. We envision that our results will have implications for the entire family of CRL proteins and their regulatory relationship with the CSN. Our study therefore provides a template not only for assisting investigations of other CRL-based systems but also for bringing together data from different structural biology techniques that otherwise will be reported independently.

## METHODS AND MATERIALS

### Preparation and expression of bacmids

WT and catalytically reduced CSN^5^H^138A^bacmids were a kind gift from Radoslav Enchev (The Francis Crick Institute, London)^19^. pcDNA3-myc3-CUL2 was a gift from Yue Xiong (Addgene plasmid #19892^33^). HA-VHL wt-pBabe-puro was a gift from William Kaelin (Addgene plasmid #19234^34^). RBX1, ELOB and ELOC were cloned from cDNA from the Mammalian Gene Collection (MGC) purchased from Dharmacon. To improve protein yield, an N-terminally truncated (1-53) natural isoform of VHL was also produced for use with mass spectrometry. Both isoforms of VHL were sub-cloned into a pET-52b(+) vector (Novagen) to add an N-terminal Step-Tag II. Genes were assembled into pACEBac1 using I-CeuI/BstXI restriction sites via the MultiBac system^35^. RBXI and CUL2 were assembled into one vector and ELOB, Strep II-VHL(ΔN) and ELOC into a second vector. Correct assembly was confirmed by sequencing of entire genes. DH10EmBacY cells were transformed with each assembly and blue/white selection was performed on L-agar plates containing 50 μg/ml kanamycin, 7 μg/ml gentamycin, 10 μg/ml tetracyclin, 100 μg/ml Bluo-Gal (Thermo Scientific) and 40 mg/ml IPTG. DH10MultiBac bacmid DNA was isolated from single white colonies. Recombinant baculoviruses were generated in Sf9 cells using standard amplification procedures.

### Expression and Purification of Recombinant CRL2

High Five Cells were co-infected with bacmids containing RBX1/CUL2 and ELOB/Strep-II VHL(ΔN)/ELOC and incubated at 27 °C and 130 rpm for 72 hours. Cells were harvested by centrifugation at 250 × g for 10 mins at 4 °C before storage at −80 °C. Freeze-thawed pellets were resuspended in 50 mM Tris pH7.5, 150 mM NaCl, 2mM DTT containing complete EDTA-free protease inhibitor tablets (Roche) and Benzonase (Sigma-Aldrich). Cells were lysed by sonication and clarified by centrifugation at 18,000 rpm for 1 hour (Beckman JA-20 rotor). Supernatant was bound to a 3 × 5 ml StrepTrap HP columns (GE Healthcare) in tandem, equilibrated with 50 mM Tris pH7.5, 150 mM NaCl, 2 mM DTT. Protein was eluted by the addition of 2.5 mM *d*-desthiobiotin. The eluted peak fractions were concentrated to 2 ml and loaded onto a Superdex 200 16/600 (GE Healthcare) size exclusion column equilibrated with 50 mM HEPES pH 7.5, 150 mM NaCl, 1 mM TCEP.

### *In vitro* Neddylation of CRL2

APPBP1-Uba3, UbcH12 and Nedd8-His were purchased from (Enzo Life Sciences). The neddylation reaction was carried out for 10 mins at 37°C with 8 μM CRL2, 350 nM APPBP1-Uba3, 1.8 μM UbcH12 and 50 μM Nedd8 in a reaction buffer (50 mM HEPES pH7.5, 150 mM NaCl) supplemented with 1.25 mM ATP and 10 mM MgCl_2_. The reaction was quenched with 15 mM DTT and ice prior to loading onto a 1 ml StrepTrap HP column (GE Healthcare). CRL2∼N8 was eluted with reaction buffer supplemented with 2.5 mM desthiobiotin and neddylation was confirmed by SDS-PAGE.

### Expression and Purification of Recombinant CSN

High Five Cells were co-infected with the bacmids gifted by Radoslav Enchev (The Francis Crick Institute, London) and protein was expressed as described for CRL2, with an additional Ni-affinity step prior to gel filtration to exploit the His6-tag on the CSN5 subunit. For Strep-affinity chromatography 50 mM HEPES pH7.5, 200 mM NaCl, 2 mM TCEP, 4% glycerol buffer was used, with the addition of 2.5 mM *d*-desthiobiotin for elution. For Ni-affinity using 2x HisTrap HP columns (GE Healthcare) in tandem, the same buffer was used, but protein was eluted by a 0-300 mM Imidazole gradient across 45 ml. For size exclusion chromatography using a Superdex200 16/600 (GE Healthcare) column was equilibrated with 50 mM HEPES pH 7.5, 150 mM NaCl, 1 mM TCEP, 2% glycerol. A second size exclusion step using a Superose 6 10/300 GL column equilibrated in 15 mM HEPES pH 7.5, 150 mM NaCl, 1 mM DTT, 1% glycerol.

### Cryo-EM of CSN-CRL2∼N8

The CSN-CRL2∼N8 complex was formed by incubation between CRL2∼N8 (1.1x molar excess) and CSN at room temperature for 90 min. The preparation (∼0.5 MDa) was subjected to size exclusion chromatography using a Superose 6 Increase 3.2/300 column (GE Healthcare), equilibrated in 15 mM HEPES pH 7.5, 100 mM NaCl, 0.5 mM DTT, 1% glycerol to reduce the contribution of apo components. Fractions from the leading edge of the peak were buffer exchanged into 15 mM HEPES pH 7.5, 100 mM NaCl using PD SpinTrap G-25 columns (GE Healthcare) before initial assessment by negative stain EM. Cryo grids were prepared using a Vitrobot (FEI). A cryo-EM dataset was collected beamline M02 from Quantifoil grids with an extra carbon layer at the Electron Bio-Imaging Centre (eBIC - Diamond Light Source, UK) on a Titan Krios 300 kV with Gatan K2 detector (M02), Å/pix = 1.06. Movies of 25 frames (dose = 1.85 e/Å^2^) were motion corrected in RELION 2.0^36^using MOTIONCOR2^37^and subsequent CTF estimation of micrographs was performed using CTFFIND4^38^(**Fig. S1**). Auto-picking selected ∼317,000 particles from ∼3100 micrographs. Particles were subjected to reference free 2D classification to assess data quality and to remove contaminants selected by auto-picking. This process reduced the particle number to ∼250,000. Following particle selection through 2D classification, particles were divided into 15 3D classes. Three of these classes (∼69,000 particles) were selected for further classification and processing, as described in **Fig. S2**.

### Cryo-EM of CSN-CRL2

The CSN-CRL2 complex was formed by incubation between CRL2 (1.1x molar excess) and CSN at room temperature for 90 min. Samples were loaded onto a 5-50% glycerol GraFix gradient containing 0-0.2% glutaraldehyde and ultracentrifuged for 24 hours at 4°C. Gradients were manually fractionated and the resultant aliquots assessed by SDS-PAGE to determine the extent of cross-linking. Fractions were also assessed using negative stain EM in-house. In order to reduce the glycerol content of samples for cryo-EM, fractions containing the desired complex (as determined by negative stain EM) were pooled together and gel filtered into 15 mM HEPES pH 7.5, 100 mM NaCl, 0.5 mM using a Superose 6 Increase 10/300 GL column (GE Healthcare). Fractions were again assessed by negative stain before preparing cryo-grids using a Vitrobot (FEI). A cryo-EM dataset was collected at the Electron Bio-Imaging Centre (eBIC - Diamond Light Source, UK) on a Titan Krios 300 kV with Gatan K2 detector (M02), with a sampling of 1.047 Å/pix. Movies of 85 frames (dose = 1.0 e/Å^2^) were motion corrected in RELION 3.0^39^using MOTIONCOR2^37^and subsequent CTF estimation of micrographs was performed using CTFFIND4^38^. Auto-picking selected ∼309,000 particles from ∼6800 micrographs. Particles were subjected to reference free 2D classification to assess data quality and to remove contaminants selected by auto-picking. This process reduced the particle number to ∼208,000. Following particle selection through 2D classification, particles were divided into 6 3D classes first with alignment, then subsequently without alignment with a mask around CSN5/CSN6 in order to perform focused classification on this area. The map that showed the greatest recovery of detail for CSN5/CSN6 (**Fig. S7**) was then subjected to 3D auto refinement and post-processing. Local resolution was estimates using ResMap^40^as part of the RELION wrapper.

### Homology modelling of the CRL2

Homology modelling of the CRL2 was necessary, due to a combination of missing domain structure for the Cullin-2 Winged-Helix A (WHA) and VHL subunit in the only crystal structure of CRL2 (PDB 5N4W). Homology modelling was performed through two stages: first, generating a CRL2 structure with a correct WHA domain, and second, generating the complete CRL2 intact with the VHL-ELOB-ELOC adaptor complex. In the first stage, we performed structural alignment of the CRL2 (5N4W) and CRL1 (1LDJ) structures in PyMOL. Using MODELLER^41^, a single model of the CRL2 was generated using the slow molecular refinement option of MODELLER. The model was manually evaluated for correct fold, including the correct positioning of residues (such as the Cullin-2 K689 NEDD8-acceptor site) already present in the 5N4W crystal structure. In the second stage, we aligned the CRL2 model with the VHL-ELOB-ELOC-Cullin-2 fragment (4WQO) to generate a template for homology modelling. The isoform 3 of VHL (missing residues 1-53) was used for modelling to maintain consistency with the experimental construct. Again, a single model of the CRL2 (with VHL/ELOB/ELOC) was generated using the slow molecular refinement option of MODELLER. The final model of the CRL2 shows a root mean squared deviation (RMSD) of 3.6 Å when compared to the initial crystal structure (5N4W) but includes a complete WHA domain and VHL subunit.

### Model fitting of EM maps

All models were fitted using CSN subunits sourced from the 4D10 crystal structure (chains A-H) and the CRL2 (VHL-ELOB-ELOC) structure generated and described in the ‘*Homology Modelling of the CRL2’* section. We performed map fitting first by performing rigid body fitting of the CSN and CRL2 subunits to each map in Chimera^42^then using the Molecular Dynamics Flexible Fitting (MDFF)^25^feature of NAMD^43^for positional refinement. In the rigid body fitting step, elongated subunits such as CSN2, CSN4 and Cullin-2 were first dissected into smaller rigid bodies to permit better fitting into their densities. Following map fitting of all CSN and CRL2 subunits, we then converted the structures into MDFF-compatible topology files using the Protein Structure File (PSF) builder function of VMD^44^. MDFF was performed in two steps: an initial energy minimisation step (scaling factor = 0.3 for 50,000 steps) which coerced each subunit into their map densities, and a second equilibration run (scaling factor = 10 for 200,000 steps) which applied molecular dynamics to produce structurally and energetically realistic structures. Secondary structure, cis-peptide and chirality characteristics of the initial models were calculated and enforced throughout each step to avoid a loss of internal structure for each subunit. For each of the CSN-CRL2∼N8 and CSN-CRL2 structures, subunits/domains which lacked clear density were not included to avoid interference with the fitting of other subunits. These were the WHB domain (Cullin-2 residues 656-745) for all maps, NEDD8 in all NEDD8-including maps, and VHL in the CSN-CRL2 map. The cross-correlation coefficient (CCC) which calculates the degree of overlap between the cryo-EM map and a simulated map of the same resolution from the atomic model, are reported for each model in **Table 1** of the manuscript.

### Native mass spectrometry

All spectral data were collected using a SYNAPT G2-Si (Waters Corp., Manchester, UK) high-definition mass spectrometer and samples were ionized using a NanoLockSpray(tm) dual electrospray inlet source (Waters Corporation). Protein samples were buffer exchanged and de-salted using Mini Bio-Spin Chromatography Columns (Bio-Rad) into pH 7.5 150 mM ammonium acetate. Native MS measurements were made at 20°C with trap collision-energy between 25-75 V.

### Hydrogen deuterium exchange mass spectrometry

Hydrogen deuterium exchange mass spectrometry (HDX-MS) experiments were performed on a Synapt G2-Si HDMS coupled to an Acquity UPLC M-Class system with HDX and automation (Waters Corporation, Manchester UK). Protein samples were prepared at a concentration of 7.5 μM. Isotope labelling was initiated by diluting 5 μl of each protein sample into 95 μl of buffer L (10 mM potassium phosphate in D_2_O pD 6.6). The protein was incubated at various time-points (0.25, 5 and 30 minutes) and then quenched in ice cold buffer Q (100 mM potassium phosphate, brought to Ph 2.3 with formic acid) before being digested online with a Waters Enzymate BEH pepsin column at 20 °C. The same procedure was used for un-deuterated control, with the labelling buffer being replaced by buffer E (10 mM potassium phosphate in H_2_O pH 7.0). The peptides were trapped on a Waters BEH C18 VanGuard pre-column for 3 minutes at a flow rate of 200 μl/min in buffer A (0.1% formic acid ∼ pH 2.5) before being applied to a Waters BEH C-18 analytical column. Peptides were eluted with a linear gradient of buffer B (8-40% gradient of 0.1% formic acid in acetonitrile) at a flow rate of 40 μl/min. All trapping and chromatography was performed at 0.5 °C to minimise back exchange. MS^E^data were acquired with a 20-30 V trap collision energy ramp for high-energy acquisition of product ions. Leucine Enkephalin (LeuEnk - Sigma) was used as a lock mass for mass accuracy correction and the mass spectrometry was calibrated with sodium iodide. The on-line Enzymate pepsin column was washed with pepsin wash (1.5 M Gu-HCl, 4% MeOH, 0.8% formic acid) recommended by the manufacturer and a blank run using the pepsin wash was performed between each sample to prevent significant peptide carry-over from the pepsin column. Optimized peptide identification and peptide coverage for all samples was performed from undeuterated controls (three-four replicates). All deuterium timepoints were performed in triplicate. Sequence identification was made from MS^E^data from the undeuterated samples using the Waters ProteinLynx Global Server 2.5.1 (PLGS). The output peptides were filtered using DynamX (v. 3.0) using the following filtering parameters: minimum intensity of 2500, minimum and maximum peptide sequence length of 5 and 30 respectively, minimum MS/MS products of 3, minimum products per amino acid of 0.1, and a minimum peptide score of 5. Additionally, all the spectra were visually examined and only those with high signal to noise ratios were used for HDX-MS analysis. The amount of relative deuterium uptake for each peptide was determined using DynamX (v. 3.0) and are not corrected for back exchange. Confidence intervals for the ΔHDX of any individual timepoint were then determined according to Houde *et al*^45^. Specifically, the 98% confidence intervals for the ΔHDX at any single timepoint were calculated per protein per timepoint.

### Chemical cross-linking mass spectrometry

20 μl of approximately 20 μM CRL2∼N8 were incubated with 1-5 mM bis(sulfosuccinimidyl)suberate (BS3) cross-linker for 1 hr at 25°C and 350 rpm in a thermomixer. After cross-linking, proteins were (i) separated by gel electrophoresis (NuPAGE) followed by in-gel digestion or (ii) digested in solution and generated peptides were pre-fractionated by gel filtration. Gel electrophoresis was performed using the NuPAGE system according to manufacturer’s protocols. In-gel digestion was performed as described before^46^. Digestion in solution was performed in the presence of RapiGest (Waters) according to manufacturer’s protocols. For gel filtration, peptides were dissolved in 30% acetonitrile (ACN), 0.1% trifluoroacetic acid (TFA) and separated on a Superdex Peptide PC 3.2/30 column (GE Healthcare) at a flowrate of 50 μl/min.

### Mass spectrometry for XL-MS

Peptides were dissolved in 2% ACN, 0.1% formic acid (FA) and separated by nano-flow liquid chromatography (Dionex UltiMate 3000 RSLC, Thermo Scientific; mobile phase A: 0.1% (v/v) FA; mobile phase B: 80% (v/v) ACN, 0.08% (v/v) FA). Peptides were loaded onto a trap column (μ-Precolumn, C18, 100 μm I.D., particle size 5 μm; Thermo Scientific) and separated with a flow rate of 300 nL/min on an analytical C18 capillary column (Acclaim PepMap 100, C18, 75 μm I.D., particle size 3 μm, 50 cm; Thermo Scientific), with a gradient of 4-90% (v/v) mobile phase B over 66 min. Separated peptides were directly eluted into a Q Exactive hybrid quadrupole-Orbitrap or an Orbitrap Fusion Tribrid Mass Spectrometer (Thermo Scientific).

Typical mass spectrometric conditions were: spray voltage of 2.3 kV; capillary temperature of 275°C; collision energy of 30%, activation Q of 0.25. The mass spectrometer was operated in data-dependent mode. Survey full scan MS spectra were acquired in the Orbitrap. The top 20 most intense ions were selected for Higher Energy Collisional Dissociation (HCD) MS/MS fragmentation in the Orbitrap. Previously selected ions within previous 30 seconds were dynamically excluded for 30 seconds. Only Ions with charge states 2-7+ were selected. Singly charged ions as well as ions with unrecognized charge state were excluded. Internal calibration of the Orbitrap was performed using the lock mass option (lock mass: m/z 445.120025)^47^.

### Data analysis for XL-MS

Raw files were converted into Mascot generic format (mgf) files using pXtract (http://pfind.ict.ac.cn/software/pXtract/index.html). Mgf’s were searched against a reduced database containing CSN and CRL2 proteins^48^. Search parameters were: instrument spectra, HCD; enzyme, trypsin; max missed cleavage sites, 3; variable modifications, oxidation (methionine) and carbamidomethylation (cysteine); cross-linker, BS3; min peptide length, 4; max peptide length, 100; min peptide mass, 400 Da; max peptide mass, 10,000 Da; False Discovery Rate, 1%. Potential cross-linked di-peptides was evaluated by their spectral quality. Circular network plots were generated using XVis^49^.

### XL-MS guided placement of the WHB, NEDD8 and VHL subunits

Cross-links determined from XL-MS for the CSN-CRL2∼N8 and CSN-CRL2 complexes were used to clarify the position of the WHB, NEDD8 and VHL subunits which lacked clear density in our cryo-EM maps. We performed XL-guided placement using the Integrative Modelling Platform (IMP)^50^, using as input, the map-fitted models of the CSN-CRL2∼N8 and CSN-CRL2. The CSN-CRL2∼N8 model post-map fitting, included all subunits except the WHB and NEDD8. Similarly, the CSN-CRL2 model included all subunits except the WHB and VHL. Separately, the subunits of each complex were initialized as coarse-grained bead models, representing each residue as a single bead. The WHB domain (Cullin-2 residues 656-745) and VHL was sourced from the homology model of the CRL2 (detailed in the section ‘Homology Modelling of the CRL2’). NEDD8 was sourced from the crystal structure of neddylated CRL5 (3DQV). WHB, NEDD8 and VHL were set as mobile rigid bodies, while all other subunits were kept stationary.

Our modelling procedure utilized two types of cross-links. The first type are pseudo-cross-links which maintain the correct topology of the complex: a single pseudo-cross-link between Cullin-2^T655^to WHB^T656^of 5 Å to mimic a covalent bond, and connections between VHL-ELOB, VHL-ELOC and VHL-Cullin-2 to maintain integrity of the VHL-ELOB-ELOC adaptor complex and its interface with Cullin-2. A single pseudo-cross-link of 10 Å was used to mimic the isopeptide bond of WHB^K689^∼N8^G76^(7.5 Å lysine side chain + ∼3 Å glycine C-terminus). The second type are cross-links determined experimentally between WHB, NEDD8 and VHL with its surrounding subunits (**Table S1-S2**) which utilized a distance threshold of 35 Å (two lysine side chains at 15 Å, BS3 linker length at 10 Å, plus 10 Å for flexibility). IMP was parametrized to perform 1000 iterations, with each iteration randomly moving WHB, NEDD8 and VHL relative to the stationary CSN and CRL subunits. IMP parameters used were num_mc_steps = 10, rb_max_trans = 2, rb_max_rot = 0.1, bead_max_trans = 0.5 and excluded volume restraint resolution = 20. The single best model was evaluated by projecting all cross-links for the complex onto the structure and confirming that all distances were below the 35 Å distance threshold. A table of cross-links can be found in Supplementary **Table S1** for the CSN-CRL2∼N8 and **Table S2** CSN-CRL2. Additionally see **Fig. S5** and **S8**.

## Supporting information

## ACKNOWLEDGEMENTS

We thank Radoslav Enchev (The Francis Crick Institute, London) for the COP9 baculovirus plasmids, and Lucie Vyletova and Ruth Knight (Institute of Cancer Research, London) for their help with baculovirus expression and insect cell maintenance. We are grateful to Chris Richardson (Institute of Cancer Research, London) for IT support and to Jane Sandall (Institute of Cancer Research, London) for laboratory support. We thank the staff at eBIC (Diamond Light Source), particularly Dan Clare and Yuriy Chaban, for their support during cryo-EM data collection. We thank Antoni Borysik (King’s College London) for helpful guidance with HDX-MS instrumentation. We thank Jurgen Claesen (Hasselt University, Belgium) for helpful interpretation of the HDX-MS data. The London Interdisciplinary Biosciences Consortium (LIDo) BBSRC Doctoral Training Partnership (BB/M009513/1) supports AML. SVF, EPM and FB are funded by Cancer Research UK (C12209/A16749). CS acknowledges funding from the Federal Ministry for Education and Research (BMBF, ZIK programme, 03Z22HN22), the European Regional Development Funds (EFRE, ZS/2016/04/78115) and the MLU Halle-Wittenberg. CM and AP are funded by the Wellcome Trust (109854/Z/15/Z) and a King’s Health Partners R&D Challenge Fund through the MRC.

## Author contributions

E.P.M. and A.P. conceived and designed the research. A.M.L. and N.C. performed molecular modelling; S.F., H.Y., F.B. and E.P.M. contributed all protein samples and EM data; Z.A., C.M. and A.P. performed native MS experiments; C.M. conducted all HDX-MS experiments; C.S. performed all XL-MS experiments; A.M.L., S.F., E.P.M. and A.P. wrote the paper with contribution from all authors.

## Data deposition

The cryo-EM density map of the holocomplex (CSN-CRL2∼N8) is deposited in the Electron Microscopy Data Bank (EMDB) under accession code EMD-XXXX. The cryo-EM density map of the substrate receptor-free (-VHL) complex is deposited under EMD-XXXX. The cryo-EM density map of the holocomplex refined around the VHL substrate receptor is deposited under EMD-XXXX. The cryo-EM density map of the complex devoid of the substrate receptor and the catalytic CSN MPN domains (-VHL/CSN5/CSN6) is deposited under EMD-XXXX. The deneddylated CSN-CRL2 map was deposited under EMD-XXXX.

## References

1 Deshaies, R. J. & Joazeiro, C. A. RING domain E3 ubiquitin ligases. Annu Rev Biochem 78, 399–434, doi:10.1146/annurev.biochem.78.101807.093809 (2009).

2 Petroski, M. D. & Deshaies, R. J. Function and regulation of cullin-RING ubiquitin ligases. Nat Rev Mol Cell Biol 6, 9–20, doi:10.1038/nrm1547 (2005).

3 Duda, D. M. et al. Structural regulation of cullin-RING ubiquitin ligase complexes. Curr Opin Struct Biol 21, 257–264, doi:10.1016/j.sbi.2011.01.003 (2011).

4 Wang, S. et al. Atlas on substrate recognition subunits of CRL2 E3 ligases. Oncotarget 7, 46707–46716, doi:10.18632/oncotarget.8732 (2016).

5 Maeda, Y. et al. CUL2 is required for the activity of hypoxia-inducible factor and vasculogenesis. J Biol Chem 283, 16084–16092, doi:10.1074/jbc.M710223200 (2008).

6 Nguyen, H. C., Yang, H., Fribourgh, J. L., Wolfe, L. S. & Xiong, Y. Insights into Cullin-RING E3 ubiquitin ligase recruitment: structure of the VHL-EloBC-Cul2 complex. Structure 23, 441–449, doi:10.1016/j.str.2014.12.014 (2015).

7 Pause, A. et al. The von Hippel-Lindau tumor-suppressor gene product forms a stable complex with human CUL-2, a member of the Cdc53 family of proteins. Proc Natl Acad Sci U S A 94, 2156–2161 (1997).

8 Cardote, T. A. F. & Ciulli, A. Structure-Guided Design of Peptides as Tools to Probe the Protein-Protein Interaction between Cullin-2 and Elongin BC Substrate Adaptor in Cullin RING E3 Ubiquitin Ligases. ChemMedChem, doi:10.1002/cmdc.201700359 (2017).

9 Soares, P. et al. Group-based optimization of potent and cell-active inhibitors of the von Hippel-Lindau (VHL) E3 ubiquitin ligase: structure-activity relationships leading to the chemical probe (2S,4R)-1-((S)-2-(1-cyanocyclopropanecarboxamido)-3,3-dimethylbutanoyl)-4-hydroxy-N-(4-(4-methylthiazol-5-yl)benzyl)pyrrolidine-2-carboxamide (VH298). J Med Chem, doi:10.1021/acs.jmedchem.7b00675 (2017).

10 Deshaies, R. J. Protein degradation: Prime time for PROTACs. Nat Chem Biol 11, 634– 635, doi:10.1038/nchembio.1887 (2015).

11 Wada, H., Yeh, E. T. & Kamitani, T. Identification of NEDD8-conjugation site in human cullin-2. Biochem Biophys Res Commun 257, 100–105, doi:10.1006/bbrc.1999.0339 (1999).

12 Scott, D. C. et al. Structure of a RING E3 trapped in action reveals ligation mechanism for the ubiquitin-like protein NEDD8. Cell 157, 1671–1684, doi:10.1016/j.cell.2014.04.037 (2014).

13 Lingaraju, G. M. et al. Crystal structure of the human COP9 signalosome. Nature 512, 161–165, doi:10.1038/nature13566 (2014).

14 Enchev, R. I., Schreiber, A., Beuron, F. & Morris, E. P. Structural insights into the COP9 signalosome and its common architecture with the 26S proteasome lid and eIF3. Structure 18, 518–527, doi:10.1016/j.str.2010.02.008 (2010).

15 Sharon, M. et al. Symmetrical modularity of the COP9 signalosome complex suggests its multifunctionality. Structure (London, England: 1993) 17, 31–40, doi:S0969-2126(08)00417-6 [pii] 10.1016/j.str.2008.10.012 [doi] (2009).

16 Enchev, R. I. et al. Structural basis for a reciprocal regulation between SCF and CSN. Cell Rep 2, 616–627, doi:10.1016/j.celrep.2012.08.019 (2012).

17 Pick, E., Hofmann, K. & Glickman, M. H. PCI complexes: Beyond the proteasome, CSN, and eIF3 Troika. Mol Cell 35, 260–264, doi:10.1016/j.molcel.2009.07.009 (2009).

18 Rozen, S. et al. CSNAP Is a Stoichiometric Subunit of the COP9 Signalosome. Cell Rep 13, 585–598, doi:10.1016/j.celrep.2015.09.021 (2015).

19 Mosadeghi, R. et al. Structural and kinetic analysis of the COP9-Signalosome activation and the cullin-RING ubiquitin ligase deneddylation cycle. Elife 5, doi:10.7554/eLife.12102 (2016).

20 Cavadini, S. et al. Cullin-RING ubiquitin E3 ligase regulation by the COP9 signalosome. Nature 531, 598–603, doi:10.1038/nature17416 (2016).

21 Cope, G. A. et al. Role of predicted metalloprotease motif of Jab1/Csn5 in cleavage of Nedd8 from Cul1. Science 298, 608–611, doi:10.1126/science.1075901 (2002).

22 Echalier, A. et al. Insights into the regulation of the human COP9 signalosome catalytic subunit, CSN5/Jab1. Proc Natl Acad Sci U S A 110, 1273–1278, doi:10.1073/pnas.1209345110 (2013).

23 Bennett, E. J., Rush, J., Gygi, S. P. & Harper, J. W. Dynamics of Cullin-RING Ubiquitin Ligase Network Revealed by Systematic Quantitative Proteomics. Cell 143, 951–965, doi:10.1016/j.cell.2010.11.017 (2010).

24 Emberley, E. D., Mosadeghi, R. & Deshaies, R. J. Deconjugation of Nedd8 from Cul1 is directly regulated by Skp1-F-box and substrate, and the COP9 signalosome inhibits deneddylated SCF by a noncatalytic mechanism. J Biol Chem 287, 29679–29689, doi:10.1074/jbc.M112.352484 (2012).

25 Trabuco, L. G., Villa, E., Schreiner, E., Harrison, C. B. & Schulten, K. Molecular dynamics flexible fitting: a practical guide to combine cryo-electron microscopy and X-ray crystallography. Methods 49, 174–180, doi:10.1016/j.ymeth.2009.04.005 (2009).

26 Cardote, T. A. F., Gadd, M. S. & Ciulli, A. Crystal Structure of the Cul2-Rbx1-EloBC-VHL Ubiquitin Ligase Complex. Structure 25, 901–911 e903, doi:10.1016/j.str.2017.04.009 (2017).

27 Rand, K. D., Zehl, M., Jensen, O. N. & Jorgensen, T. J. Protein hydrogen exchange measured at single-residue resolution by electron transfer dissociation mass spectrometry. Anal Chem 81, 5577–5584, doi:10.1021/ac9008447 (2009).

28 Reading, E. et al. Interrogating membrane protein conformational dynamics within native lipid compositions. Angew Chem Int Ed Engl, doi:10.1002/anie.201709657 (2017).

29 Konermann, L., Pan, J. & Liu, Y. H. Hydrogen exchange mass spectrometry for studying protein structure and dynamics. Chem Soc Rev 40, 1224–1234, doi:10.1039/c0cs00113a (2011).

30 Wales, T. E. & Engen, J. R. Hydrogen exchange mass spectrometry for the analysis of protein dynamics. Mass spectrometry reviews 25, 158–170, doi:10.1002/mas.20064 (2006).

31 Mistarz, U. H., Brown, J. M., Haselmann, K. F. & Rand, K. D. Probing the Binding Interfaces of Protein Complexes Using Gas-Phase H/D Exchange Mass Spectrometry. Structure (London, England: 1993) 24, 310–318, doi:10.1016/j.str.2015.11.013 (2016).

32 Marcsisin, S. R. & Engen, J. R. Hydrogen exchange mass spectrometry: what is it and what can it tell us? Analytical and bioanalytical chemistry 397, 967–972, doi:10.1007/s00216-010-3556-4 (2010).

33 Ohta, T., Michel, J. J., Schottelius, A. J. & Xiong, Y. ROC1, a homolog of APC11, represents a family of cullin partners with an associated ubiquitin ligase activity. Mol Cell 3, 535–541 (1999).

34 Li, L. et al. Hypoxia-inducible factor linked to differential kidney cancer risk seen with type 2A and type 2B VHL mutations. Mol Cell Biol 27, 5381–5392, doi:10.1128/MCB.00282-07 (2007).

35 Sari, D. et al. The MultiBac Baculovirus/Insect Cell Expression Vector System for Producing Complex Protein Biologics. Adv Exp Med Biol 896, 199–215, doi:10.1007/978-3-319-27216-0_13 (2016).

36 Scheres, S. H. RELION: implementation of a Bayesian approach to cryo-EM structure determination. Journal of structural biology 180, 519–530, doi:10.1016/j.jsb.2012.09.006 (2012).

37 Zheng, S. Q. et al. MotionCor2: anisotropic correction of beam-induced motion for improved cryo-electron microscopy. Nat Methods 14, 331–332, doi:10.1038/nmeth.4193 (2017).

38 Rohou, A. & Grigorieff, N. CTFFIND4: Fast and accurate defocus estimation from electron micrographs. J Struct Biol 192, 216–221, doi:10.1016/j.jsb.2015.08.008 (2015).

39 Zivanov, J. et al. New tools for automated high-resolution cryo-EM structure determination in RELION-3. Elife 7, doi:10.7554/eLife.42166 (2018).

40 Kucukelbir, A., Sigworth, F. J. & Tagare, H. D. Quantifying the local resolution of cryo-EM density maps. Nat Methods 11, 63–65, doi:10.1038/nmeth.2727 (2014).

41 Webb, B. & Sali, A. Comparative Protein Structure Modeling Using MODELLER. Curr Protoc Protein Sci 86, 2 9 1–2 9 37, doi:10.1002/cpps.20 (2016).

42 Pettersen, E. F. et al. UCSF Chimera--a visualization system for exploratory research and analysis. J Comput Chem 25, 1605–1612, doi:10.1002/jcc.20084 (2004).

43 Phillips, J. C. et al. Scalable molecular dynamics with NAMD. J Comput Chem 26, 1781– 1802, doi:10.1002/jcc.20289 (2005).

44 Humphrey, W., Dalke, A. & Schulten, K. VMD: visual molecular dynamics. J Mol Graph 14, 33–38, 27-38 (1996).

45 Houde, D., Berkowitz, S. A. & Engen, J. R. The utility of hydrogen/deuterium exchange mass spectrometry in biopharmaceutical comparability studies. J Pharm Sci 100, 2071– 2086, doi:10.1002/jps.22432 (2011).

46 Shevchenko, A., Wilm, M., Vorm, O. & Mann, M. Mass spectrometric sequencing of proteins silver-stained polyacrylamide gels. Anal Chem 68, 850–858 (1996).

47 Olsen, J. V. et al. Parts per million mass accuracy on an Orbitrap mass spectrometer via lock mass injection into a C-trap. Mol Cell Proteomics 4, 2010–2021, doi:10.1074/mcp.T500030-MCP200 (2005).

48 Yang, B. et al. Identification of cross-linked peptides from complex samples. Nat Methods 9, 904–906, doi:10.1038/nmeth.2099 (2012).

49 Grimm, M., Zimniak, T., Kahraman, A. & Herzog, F. xVis: a web server for the schematic visualization and interpretation of crosslink-derived spatial restraints. Nucleic Acids Res 43, W362–369, doi:10.1093/nar/gkv463 (2015).

50 Russel, D. et al. Putting the pieces together: integrative modeling platform software for structure determination of macromolecular assemblies. PLoS Biol 10, e1001244, doi:10.1371/journal.pbio.1001244 (2012).

