## Supplementary material for "Structural basis of Cullin-2 RING E3 ligase regulation by the COP9 signalosome"

**This PDF file includes:**

Supplementary Figures (Fig. S1 to S15)

Supplementary Tables (Table S1 to S2)

Supplementary References

#### Supplementary Figures

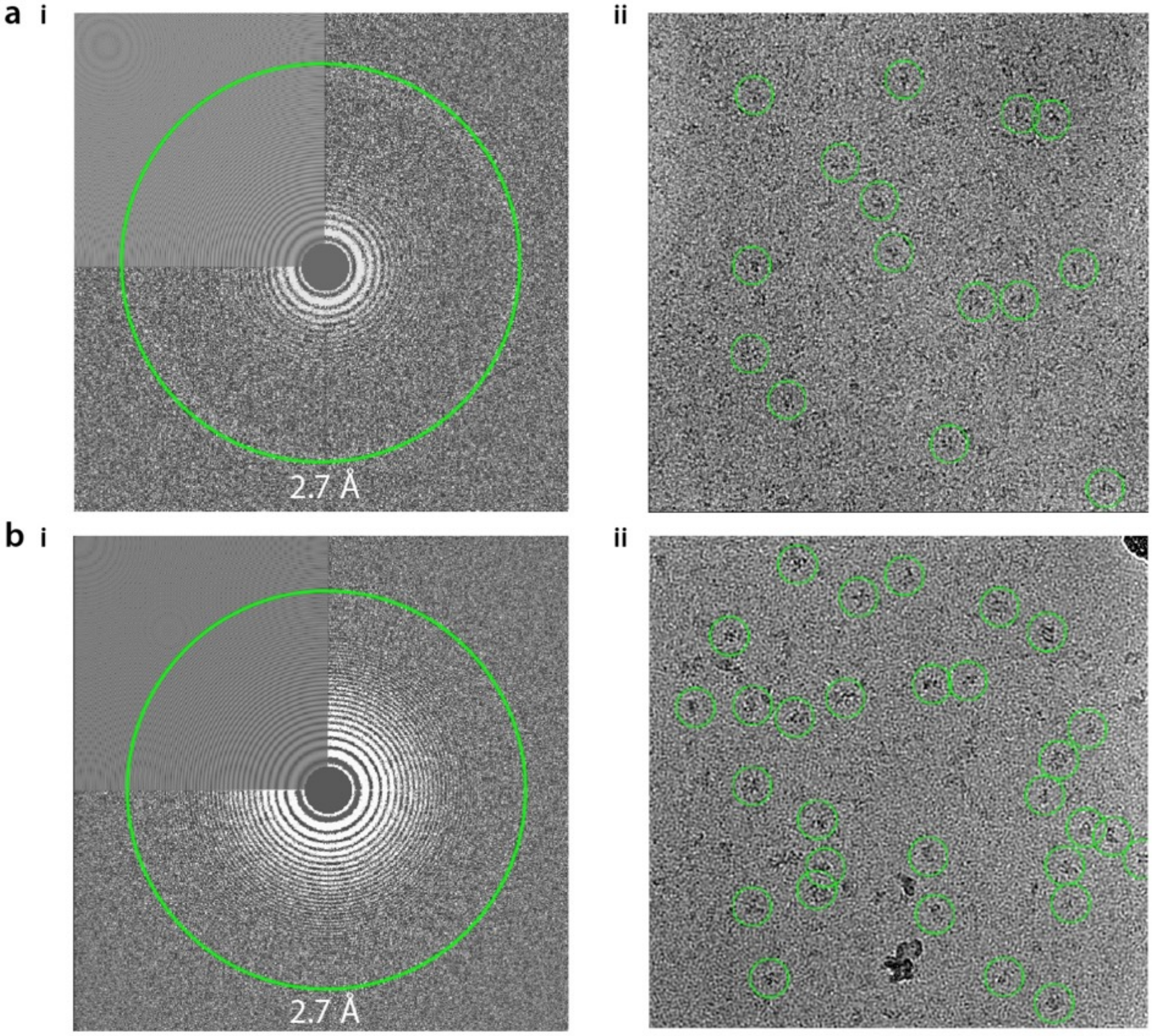


**Fig. S1. Cryo-electron micrographs.** Micrographs of (**a**) CSN-CRL2~N8 and (**b**) CSN-CRL2 complexes. (**i**) Representative power spectrum and (**ii**) various single molecular views are circled.

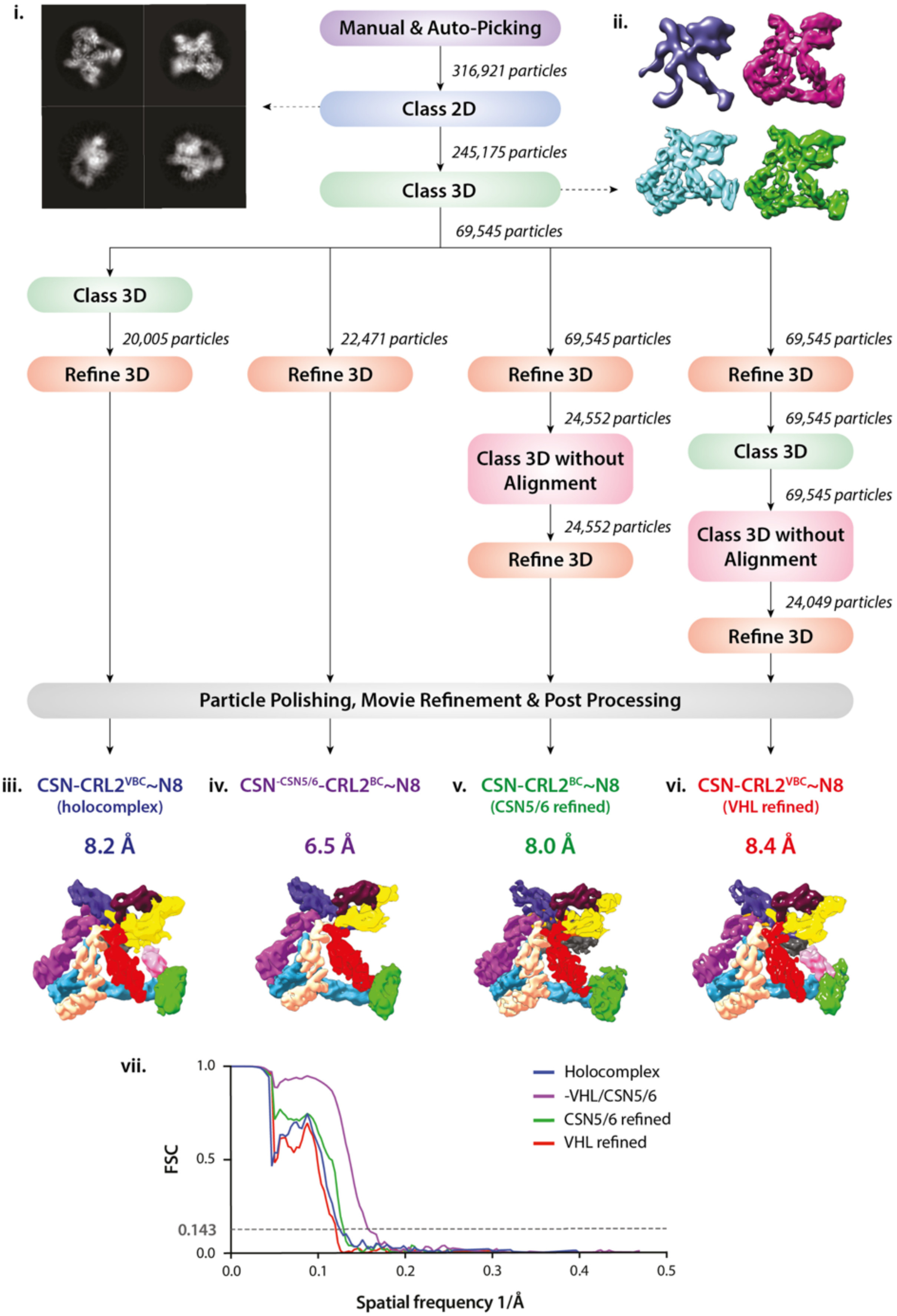
i.
ii.
Manual & Auto-Picking
316,921 particles
Class 2D
245,175 particles
Class 3D
69,545 particles
Class 3D
20,005 particles
Refine 3D
22,471 particles
Refine 3D
69,545 particles
Refine 3D
24,552 particles
Class 3D without Alignment
24,552 particles
Refine 3D
69,545 particles
Refine 3D
69,545 particles
Class 3D
69,545 particles
Class 3D without Alignment
24,049 particles
Refine 3D
Particle Polishing, Movie Refinement & Post Processing
iii. CSN-CRL2VBC~N8 (holocomplex)
8.2 Å
iv. CSN-CSN5/6-CRL2BC~N8
6.5 Å
v. CSN-CRL2BC~N8 (CSN5/6 refined)
8.0 Å
vi. CSN-CRL2VBC~N8 (VHL refined)
8.4 Å
vii.
FSC
1.0
0.5
0.143
0.0
0.0
0.1
0.2
0.3
0.4
0.5
Spatial frequency 1/Å
Holocomplex
-VHL/CSN5/6
CSN5/6 refined
VHL refined

**Fig. S2. Flowchart depicting the workflow for processing cryo-EM data**. A set of ~3100 micrographs were subjected to manual and auto-picking in order to acquire particles for 2D reference-free classification (**i**). 2D classification was used for the positive selection of particles prior to 3D classification. (**ii**) Four of the fifteen classes generated, demonstrate subunit heterogeneity and presence of a small component of apo-CSN (purple model) in the data set. Particles from the three models containing the CSN and CRL2~N8 in (**ii**) were pooled and further processed as described in the workflow. Maps (**v**) (CSN-CRL2$^{-VHL}$~N8) and (**vi**) (CSN-CRL2 ~N8) were generated using focused refinement by applying a mask and using the area of interest as a starting model. (**vii**) shows the FSCs for the four maps (**iii-vi**).

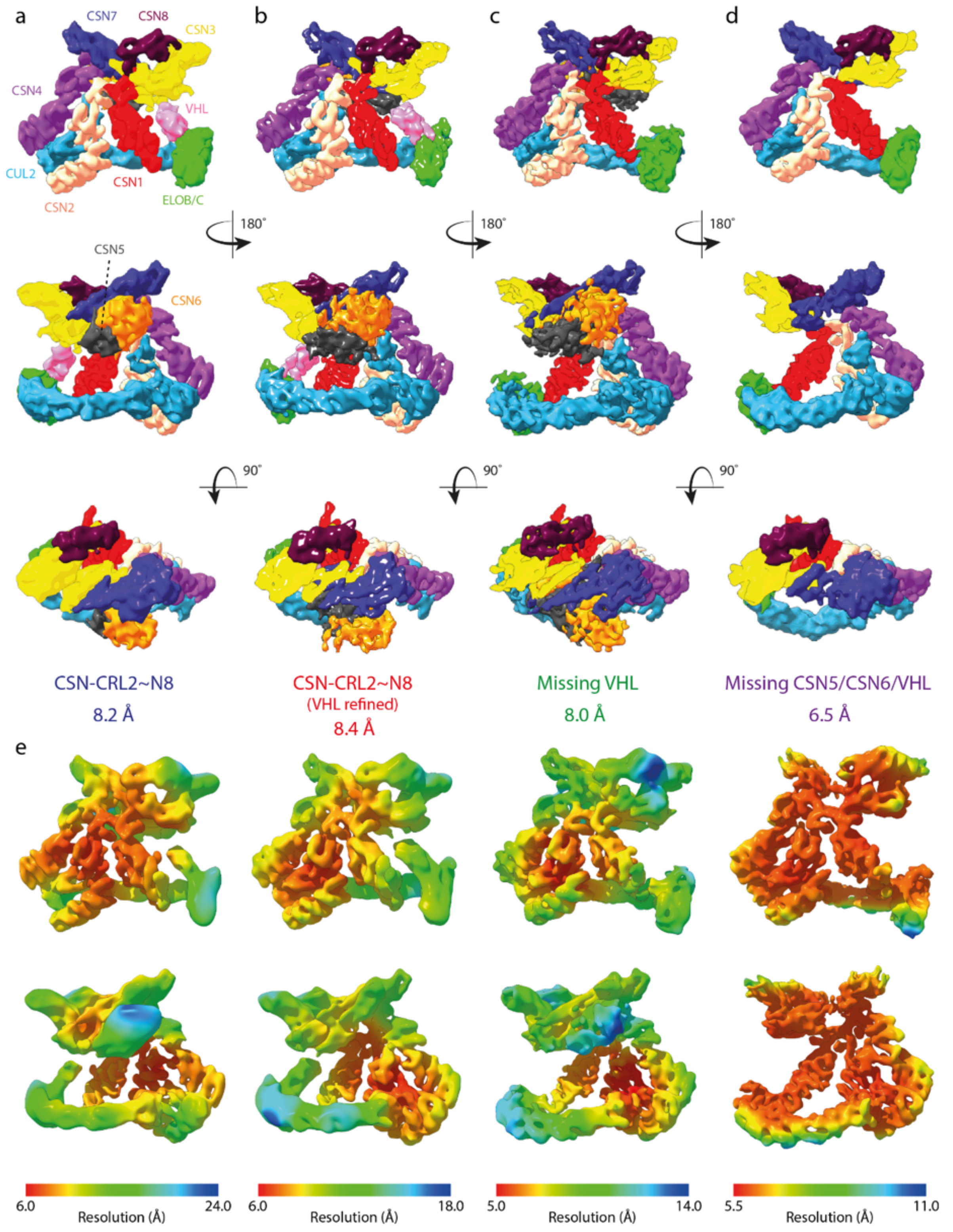


**Fig. S3. Cryo-EM structures of the CSN-CRL2~N8 segmented to highlight subunit composition.** Maps of the (**a**) CSN-CRL2~N8 holocomplex, (**b**) holocomplex (from VHL focused refinement), (**c**) holocomplex missing VHL (**d**) holocomplex missing CSN5/CSN6 and

VHL, are shown in various orientations. (**e**) Resolution maps of (**a-d**) from front and back views. Map resolutions generated using RELION[1] software.

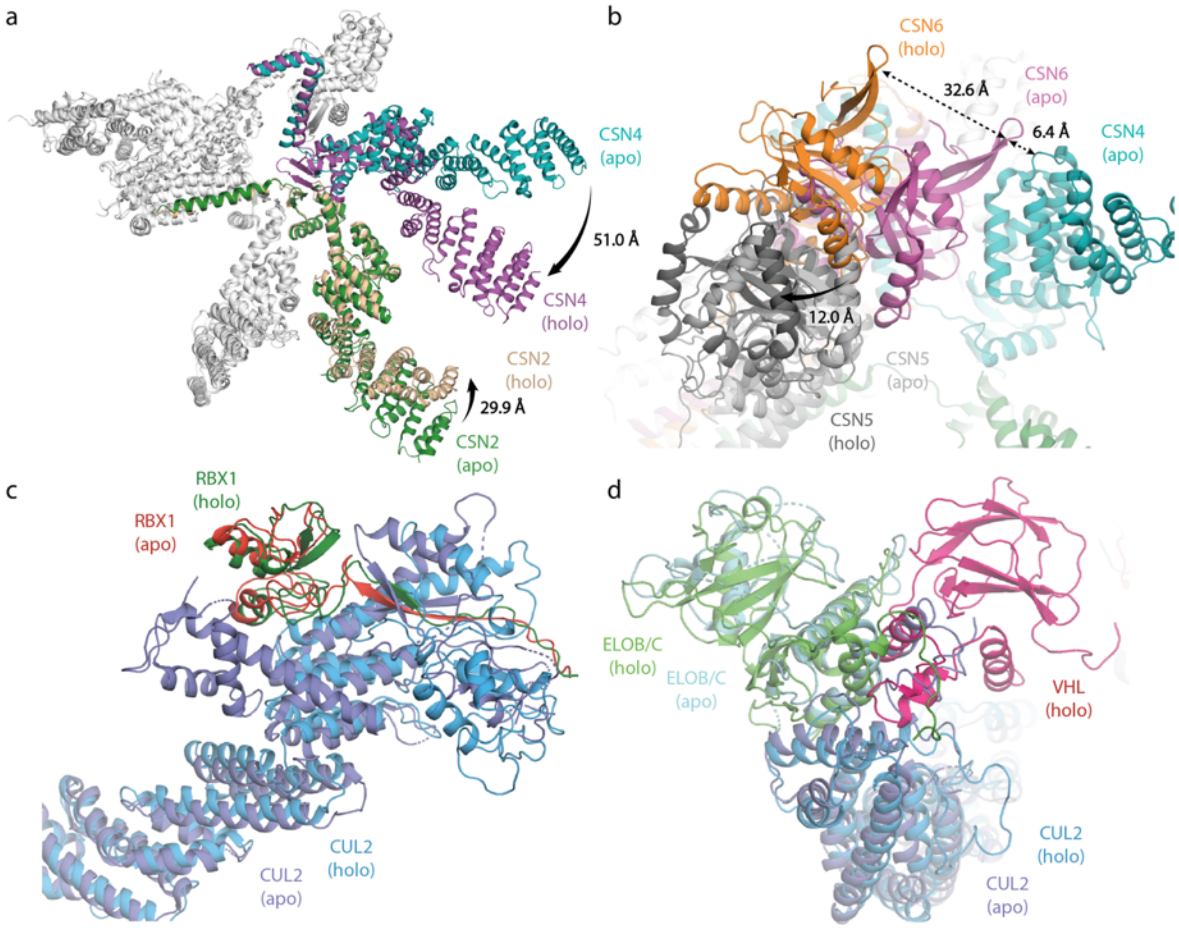


**Fig. S4. Comparisons of the apo-CSN and CSN-CRL2~N8 structures.** Large scale conformational changes of (**a**) CSN2/CSN4 and (**b**) CSN5/CSN6 upon binding of CRL2~N8 (holo). Subunits of the CSN-CRL2~N8 were compared to the apo-CSN crystal structure (PDB 4D10) following structural alignment. The structure of the CRL2~N8 has been hidden for clarity. Comparisons of the CRL2 (**c**) N-terminal domain and (**d**) C-terminal domain in isolated (apo; 5N4W) and CSN-associated (holo) conformations.

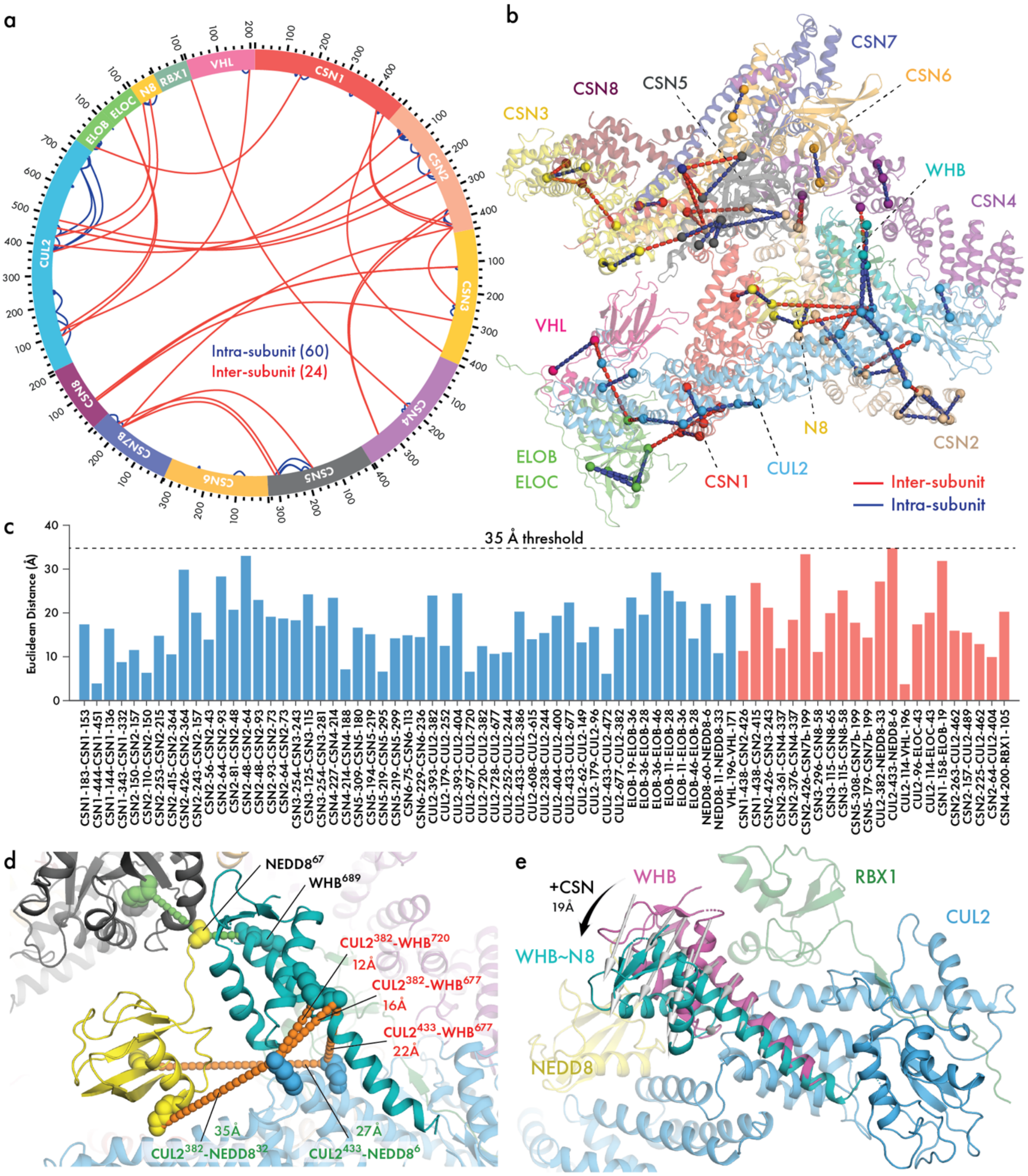


**Fig. S5. Cross-links of the CSN-CRL2~N8 complex.** (**a**) Circular plot of cross-links identified for the CSN-CRL2~N8 complex. (**b**) Inter- (red) and intra-subunit (blue) cross-links projected onto the cryo-EM structure of CSN-CRL2~N8. Nine cross-links were omitted due to missing residues in the C-terminal loops of CSN subunits (six) and self-residue cross-links (three) (**Table S1**). (**c**) Euclidean distances for cross-links shown in (**b**). All measurements made between lysine Nζ-Nζ atoms using in house scripts. 35 Å cross-link distance threshold is shown by the dotted line. 35 Å takes into account two lysine sidechains (15 Å), 10 Å cross-linker length for BS3 and an additional 10 Å to account for domain-level flexibility. (**d**) Zoom of the WHB~N8 region of the CSN-CRL2~N8 model generated from cryo-EM and cross-link. The WHB domain is shown in teal, CUL2 in light blue (transparent), NEDD8 in yellow and CSN5 in gray. (**e**) Comparison of WHB~N8 from our hybrid cryo-EM/cross-linking CSN-CRL2~N8 model (teal) with non-neddylated WHB from CRL2 crystal structure (5N4W; purple). Vector arrows depict the 19 Å movement of WHB following neddylation and finding to CSN.

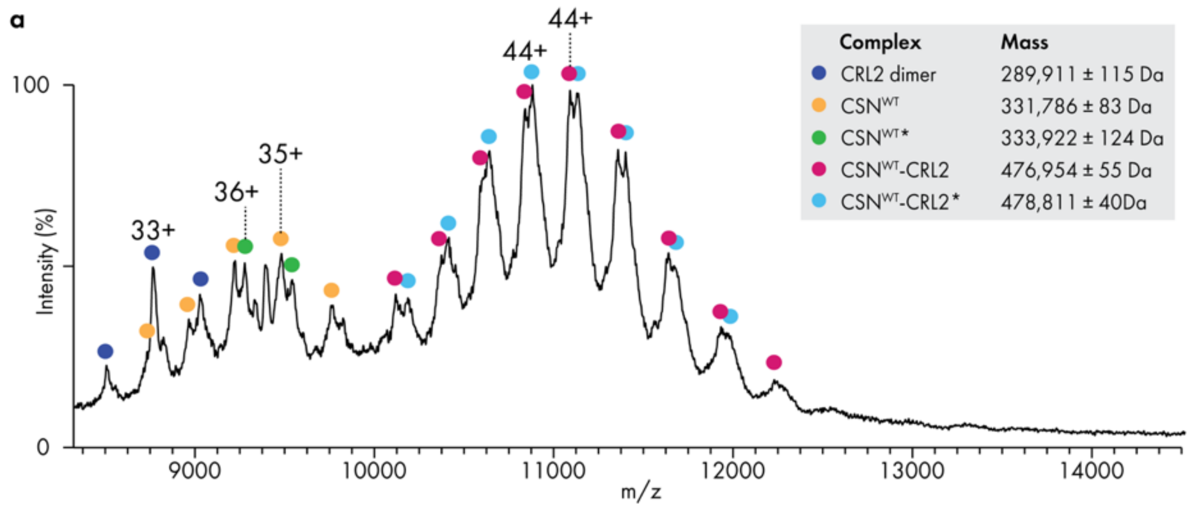


**Fig. S6. Native MS of the CSN$^{WT}$-CRL2.** Mass were assigned using Waters MassLynx software. Complex assignment marked by the asterisk (*) indicate masses which show an unidentified additional ~2 kDa mass increase.

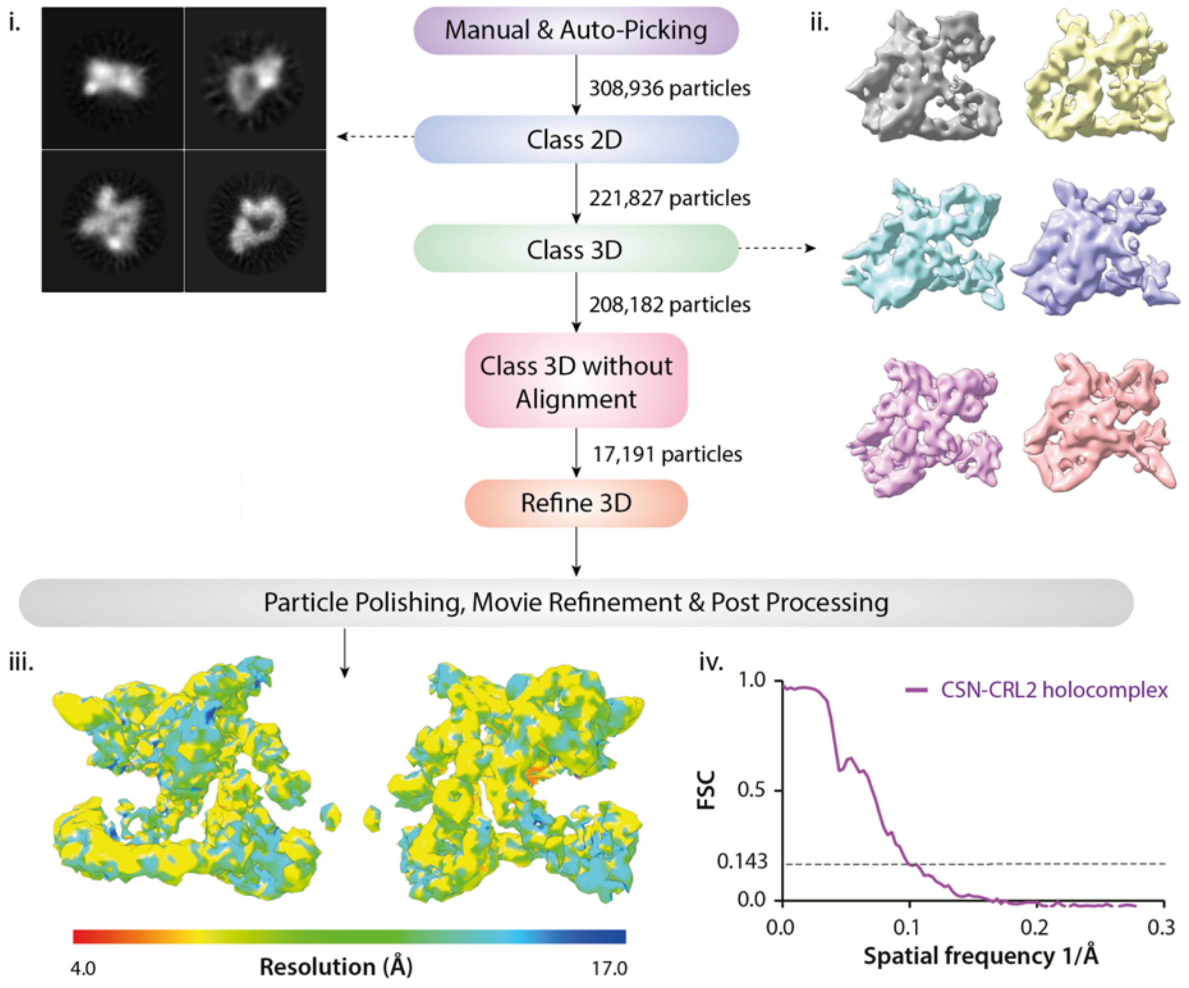


**Fig. S7. Cryo-EM map of the CSN-CRL2 complex.** A set of 6800 micrographs were subjected to manual and auto-picking in order to acquire particles for 2D reference-free classification (**i**). 2D classification was used for the positive selection of particles prior to 3D classification. (**ii**) six classes generated, demonstrate subunit heterogeneity in the data set. (**iii**) Resolution map of the CSN-CRL2 was generated using RELION[1]. (**iv**) FSCs for the single CSN-CRL2 map.

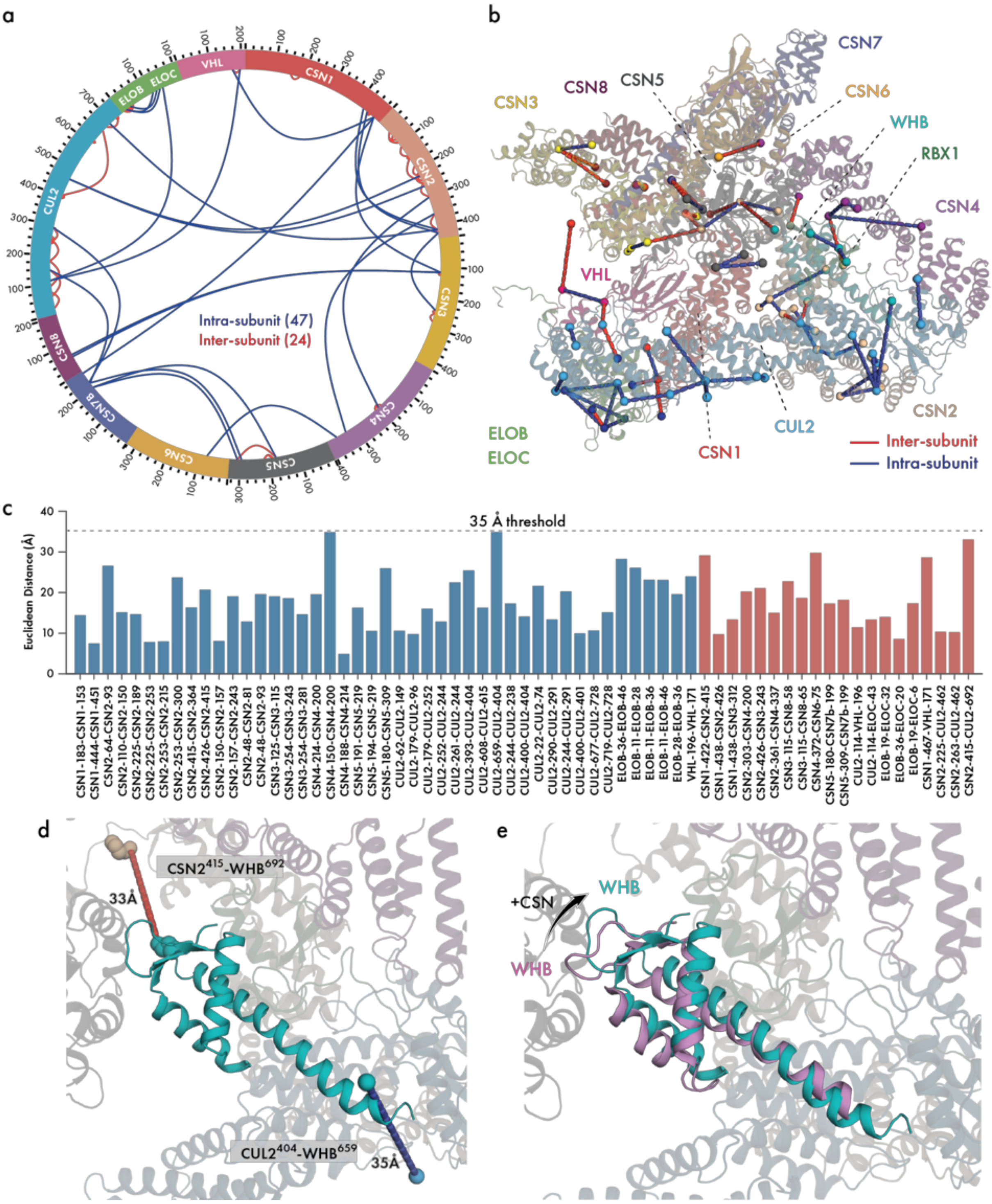


**Fig. S8. Cross-links of the deneddylated CSN-CRL2 complex.** (**a**) Circular plot of cross-links identified for the CSN-CRL2 complex. (**b**) Inter- (red) and intra-subunit (blue) cross-links projected onto the cryo-EM structure of CSN-CRL2. Six cross-links were omitted due to missing residues in the C-terminal loops of CSN subunits (five) and self-residue cross-links (one) (**Table S2**). (**c**) Euclidean distances for cross-links shown in (**b**). All measurements made between lysine Nζ-Nζ atoms using in house scripts. 35 Å cross-link distance threshold is shown by the dotted line. 35 Å takes into account two lysine sidechains (15 Å), 10 Å cross-linker length for BS3 and an additional 10 Å to account for domain-level flexibility. (**e**) Comparison of deneddylated WHB from our hybrid cryo-EM/cross-linking CSN-CRL2 model (teal) with WHB from CRL2 crystal structure (5N4W; purple).

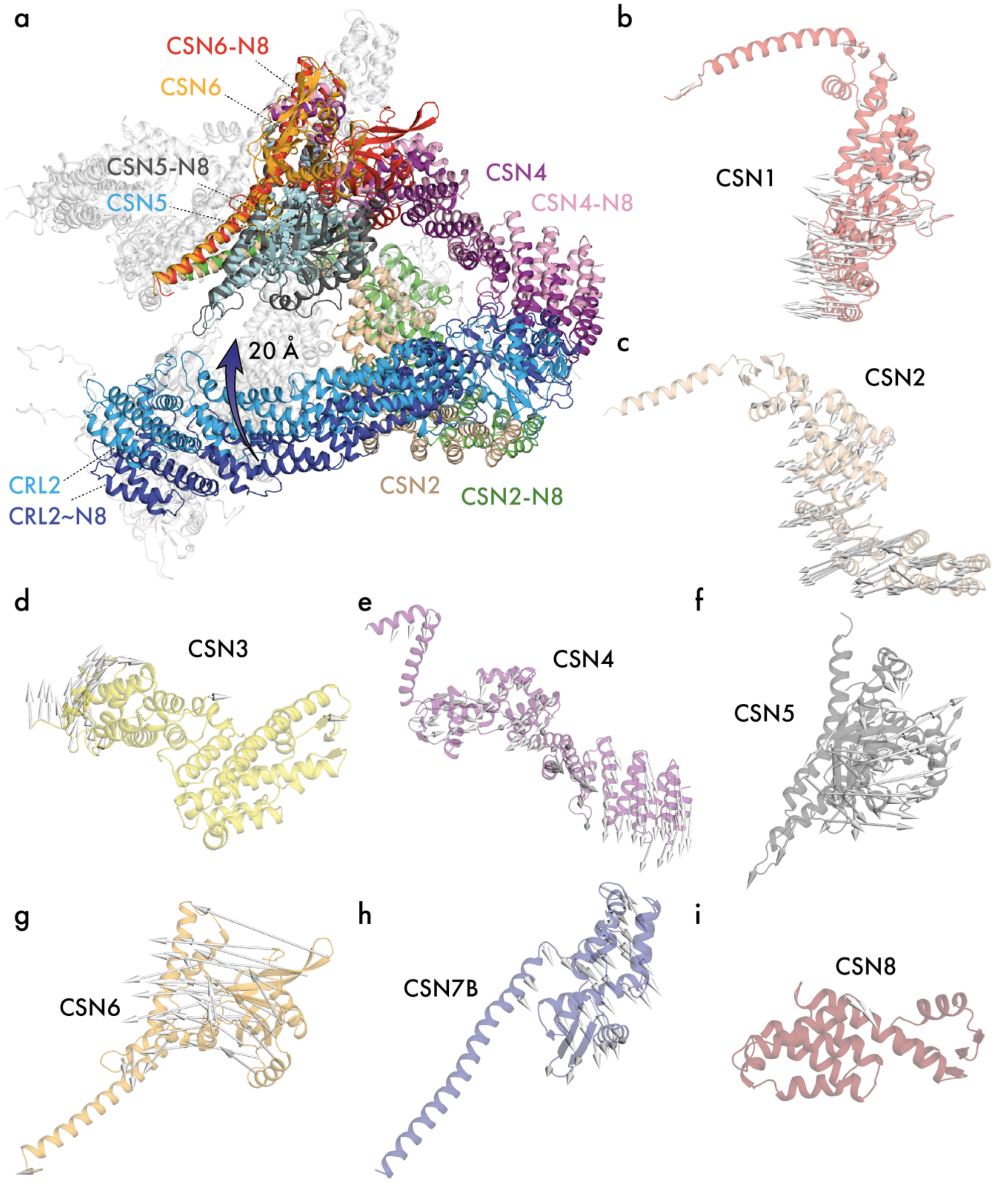


**Fig. S9. Per-subunit comparisons of the CSN in neddylated and non-neddylated CSN-CRL2 complexes.** (**a**) Superposition of the CSN-CRL2~N8 and CSN-CRL2. CSN1, CSN3, CSN7B, CSN8, RBX1, ELOB/C and VHL are shown in white to highlight the changes in CSN5/CSN6, CSN2/CSN4 and Cullin-2. Cullin-2 rotates upwards by ~20 Å in the absence of NEDD8. Per-subunit comparisons of (**b**) CSN1, (**c**) CSN2, (**d**) CSN3, (**e**), CSN4, (**f**) CSN5, (**g**) CSN6, (**h**) CSN7B, (**i**) CSN8 between CSN-CRL2~N8 and CSN-CRL2 structures. Areas of change have been displayed as vectors (white arrows). Vectors are plotted onto the subunits of CSN-CRL2~N8 in the direction of change from CSN-CRL2~N8 to CSN-CRL2.

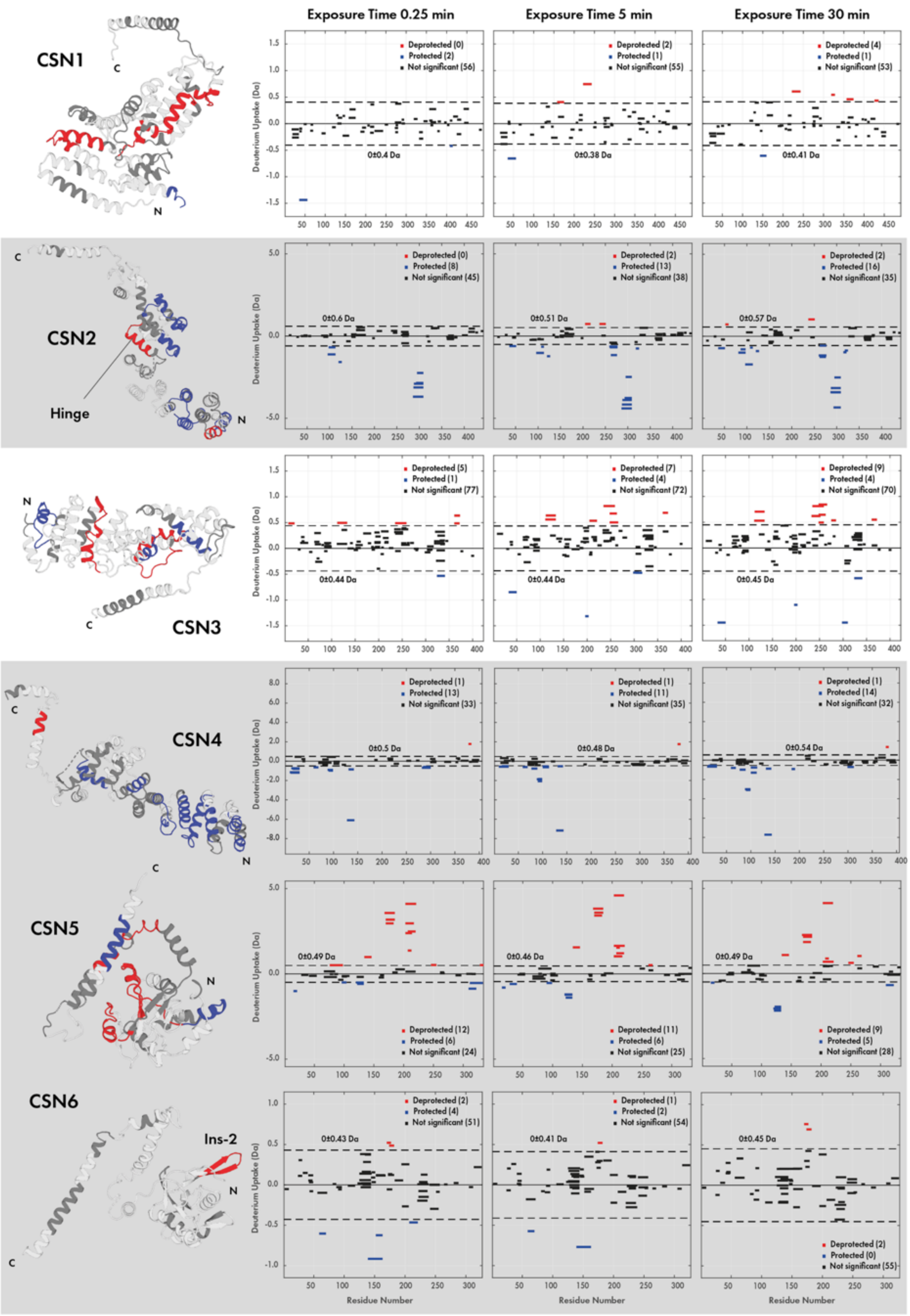
Exposure Time 0.25 min
Exposure Time 5 min
Exposure Time 30 min
CSN1
CSN2
Hinge
CSN3
CSN4
CSN5
CSN6
Ins-2
Deuterium Uptake (Da)
Residue Number
Deprotected (0)
Protected (2)
Not significant (56)
0±0.4 Da
Deprotected (2)
Protected (1)
Not significant (55)
0±0.38 Da
Deprotected (4)
Protected (1)
Not significant (53)
0±0.41 Da
Deprotected (0)
Protected (8)
Not significant (45)
0±0.6 Da
Deprotected (2)
Protected (13)
Not significant (38)
0±0.51 Da
Deprotected (2)
Protected (16)
Not significant (35)
0±0.57 Da
Deprotected (5)
Protected (1)
Not significant (77)
0±0.44 Da
Deprotected (7)
Protected (4)
Not significant (72)
0±0.44 Da
Deprotected (9)
Protected (4)
Not significant (70)
0±0.45 Da
Deprotected (1)
Protected (13)
Not significant (33)
0±0.5 Da
Deprotected (1)
Protected (11)
Not significant (35)
0±0.48 Da
Deprotected (1)
Protected (14)
Not significant (32)
0±0.54 Da
Deprotected (12)
Protected (6)
Not significant (24)
0±0.49 Da
Deprotected (11)
Protected (6)
Not significant (25)
0±0.46 Da
Deprotected (9)
Protected (5)
Not significant (28)
0±0.49 Da
Deprotected (2)
Protected (4)
Not significant (51)
0±0.43 Da
Deprotected (1)
Protected (2)
Not significant (54)
0±0.41 Da
Deprotected (2)
Protected (0)
Not significant (55)
0±0.45 Da

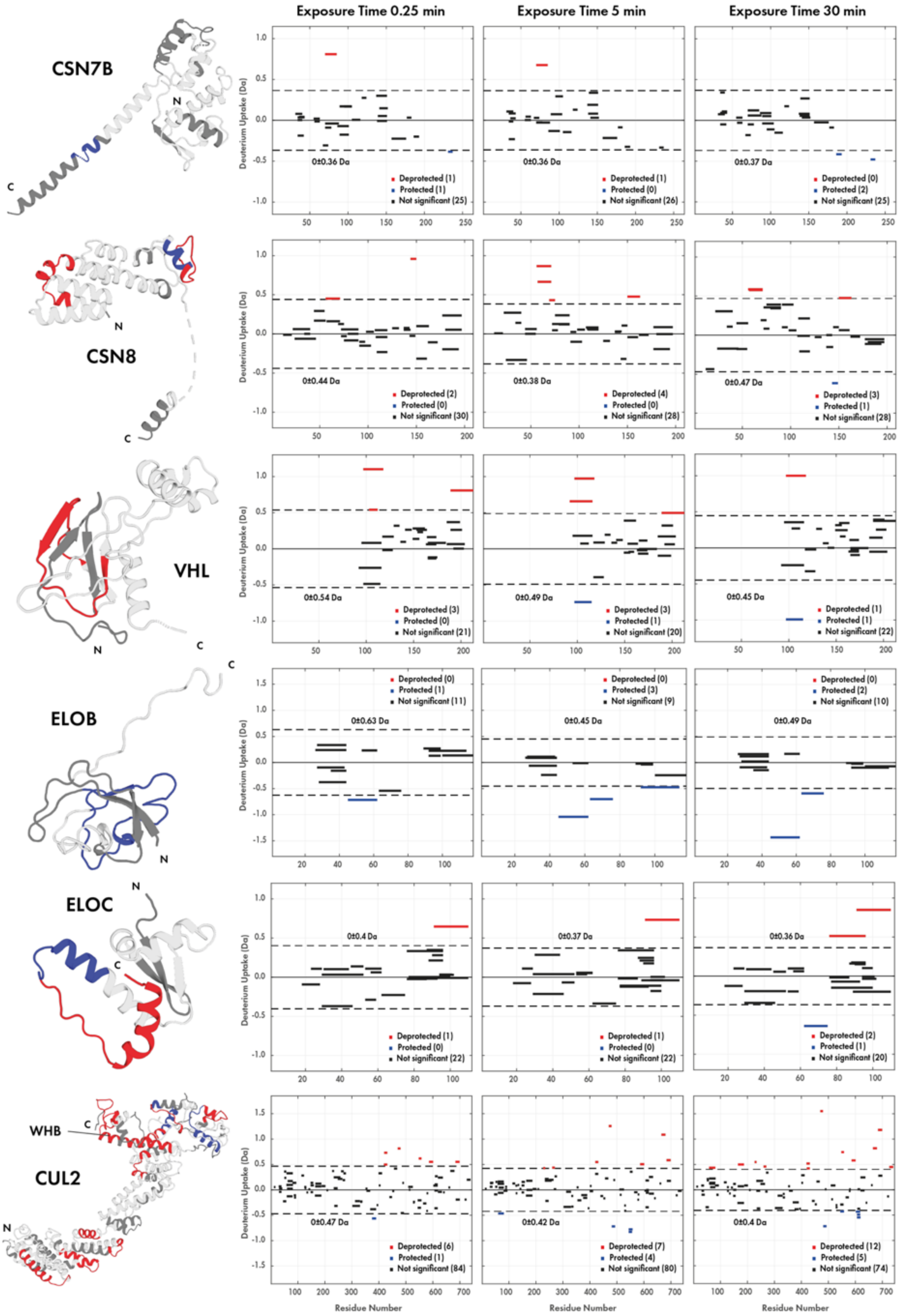
Exposure Time 0.25 min
Exposure Time 5 min
Exposure Time 30 min
CSN7B
CSN8
VHL
ELOB
ELOC
WHB
CUL2
Deuterium Uptake (Da)
Residue Number
Deprotected
Protected
Not significant

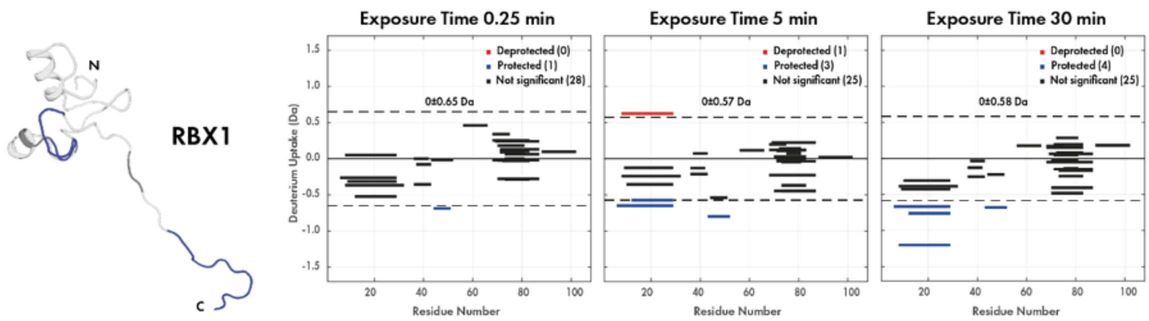


**Fig. S10. HDX-MS of CSN-CRL2~N8 per protein per timepoint.** Differential comparison of Δ(CSN-CRL2~N8 – CSN). Peptides experiencing stabilisation upon CRL2~N8 binding to CSN, compared to the peptide in apo CSN, are shown as blue, destabilised peptides are shown in red. CSN2, CSN4, CSN5 and CSN6 have been highlighted by gray boxes. The CSN2 hinge and Cullin-2 WHB domain have been highlighted in grey for clarity. Peptides were filtered applying a 98% confidence limit (critical value of 6.965; dotted lines). Structures show data for the 30 min timepoint.

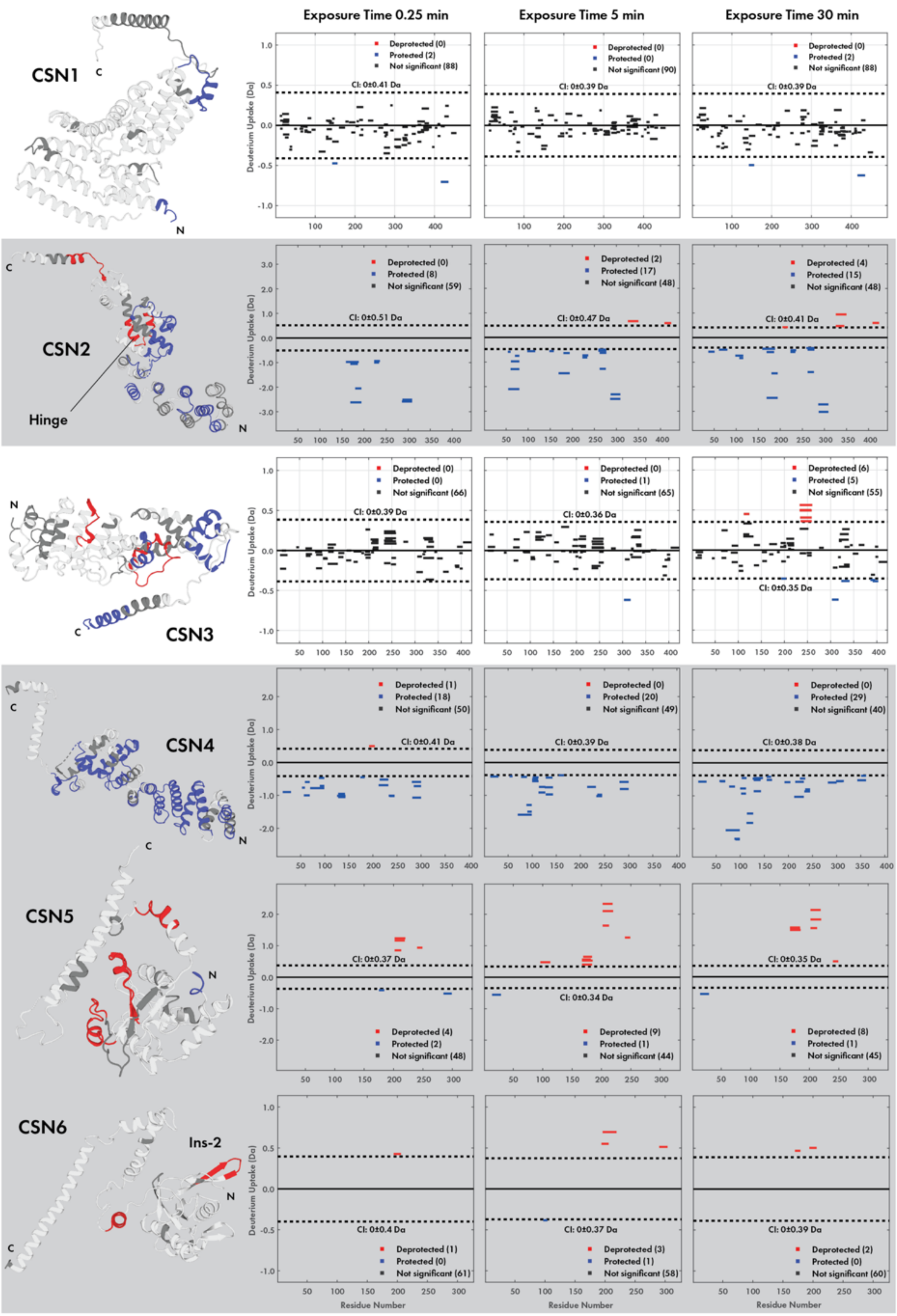

Exposure Time 0.25 min
Exposure Time 5 min
Exposure Time 30 min
CSN1
CSN2
Hinge
CSN3
CSN4
CSN5
CSN6
Ins-2
Deuterium Uptake (Da)
Residue Number
Deprotected
Protected
Not significant

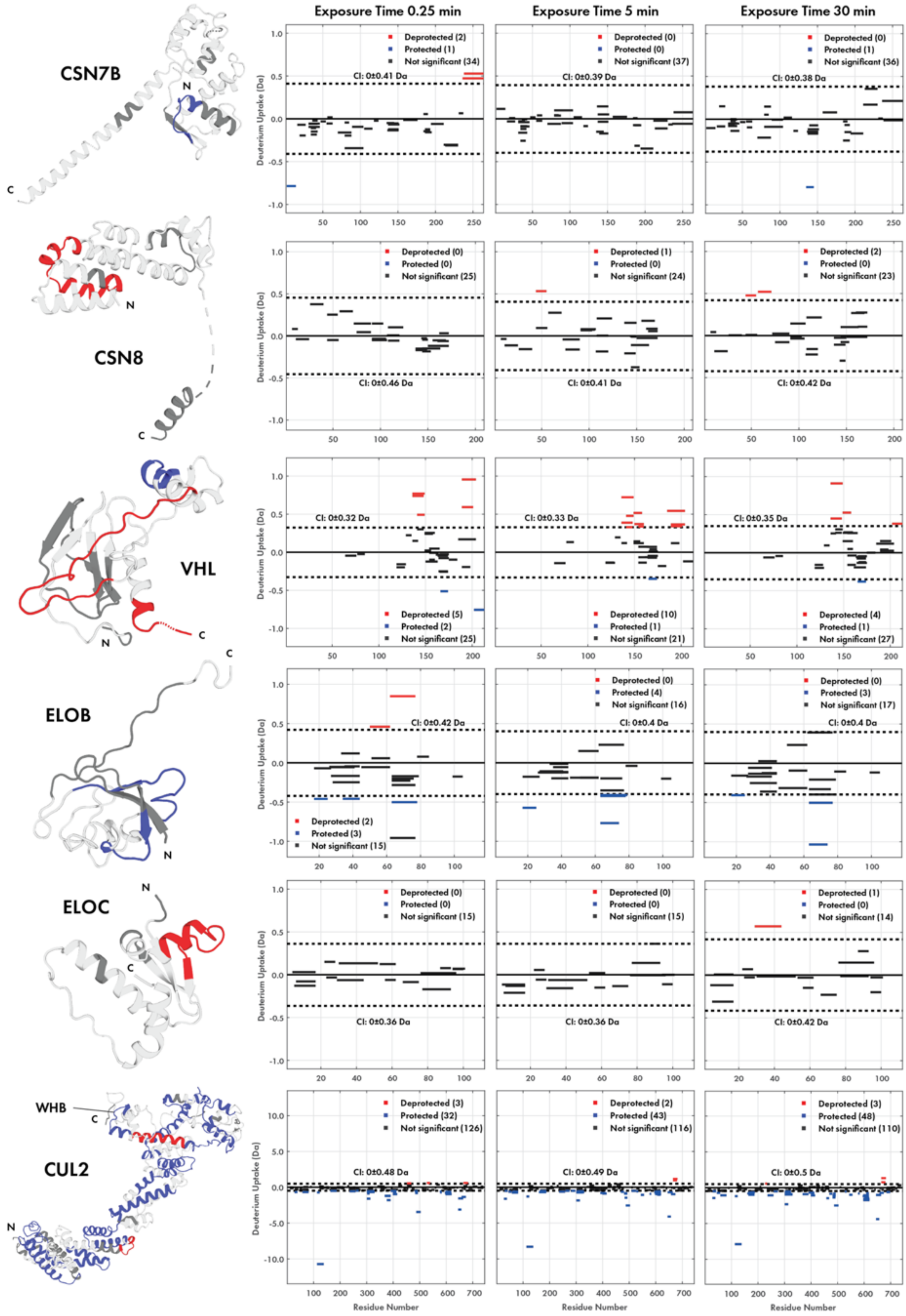

Exposure Time 0.25 min
Exposure Time 5 min
Exposure Time 30 min
CSN7B
Deprotected (2)
Protected (1)
Not significant (34)
CI: 0±0.41 Da
Deprotected (0)
Protected (0)
Not significant (37)
CI: 0±0.39 Da
Deprotected (0)
Protected (1)
Not significant (36)
CI: 0±0.38 Da
CSN8
Deprotected (0)
Protected (0)
Not significant (25)
CI: 0±0.46 Da
Deprotected (1)
Protected (0)
Not significant (24)
CI: 0±0.41 Da
Deprotected (2)
Protected (0)
Not significant (23)
CI: 0±0.42 Da
VHL
CI: 0±0.32 Da
Deprotected (5)
Protected (2)
Not significant (25)
CI: 0±0.33 Da
Deprotected (10)
Protected (1)
Not significant (21)
CI: 0±0.35 Da
Deprotected (4)
Protected (1)
Not significant (27)
ELOB
CI: 0±0.42 Da
Deprotected (2)
Protected (3)
Not significant (15)
Deprotected (0)
Protected (4)
Not significant (16)
CI: 0±0.4 Da
Deprotected (0)
Protected (3)
Not significant (17)
CI: 0±0.4 Da
ELOC
Deprotected (0)
Protected (0)
Not significant (15)
CI: 0±0.36 Da
Deprotected (0)
Protected (0)
Not significant (15)
CI: 0±0.36 Da
Deprotected (1)
Protected (0)
Not significant (14)
CI: 0±0.42 Da
WHB
CUL2
Deprotected (3)
Protected (32)
Not significant (126)
CI: 0±0.48 Da
Deprotected (2)
Protected (43)
Not significant (116)
CI: 0±0.49 Da
Deprotected (3)
Protected (48)
Not significant (110)
CI: 0±0.5 Da
Deuterium Uptake (Da)
Residue Number

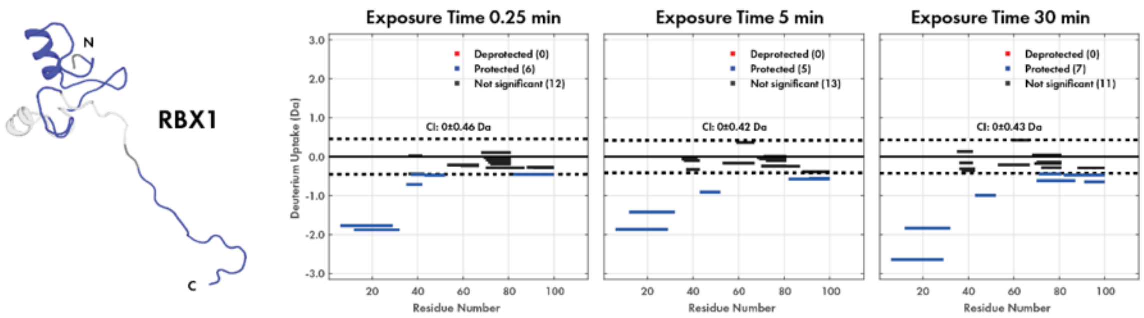


**Fig. S11. HDX-MS of CSN$^{WT}$-CRL2 per protein per timepoint.** Differential comparison of Δ(CSN$^{WT}$-CRL2 – CSN$^{WT}$). Peptides experiencing stabilisation upon non-neddylated CRL2 binding to CSN$^{WT}$, compared to the peptide in apo CSN$^{WT}$, are shown as blue, destabilised peptides are shown in red. CSN2, CSN4, CSN5 and CSN6 have been highlighted by gray boxes. The CSN2 hinge and Cullin-2 WHB domain have been highlighted for clarity. Peptides were filtered applying a 98% confidence limit (critical value of 6.965; dotted lines). Structures show data for the 30 min timepoint.

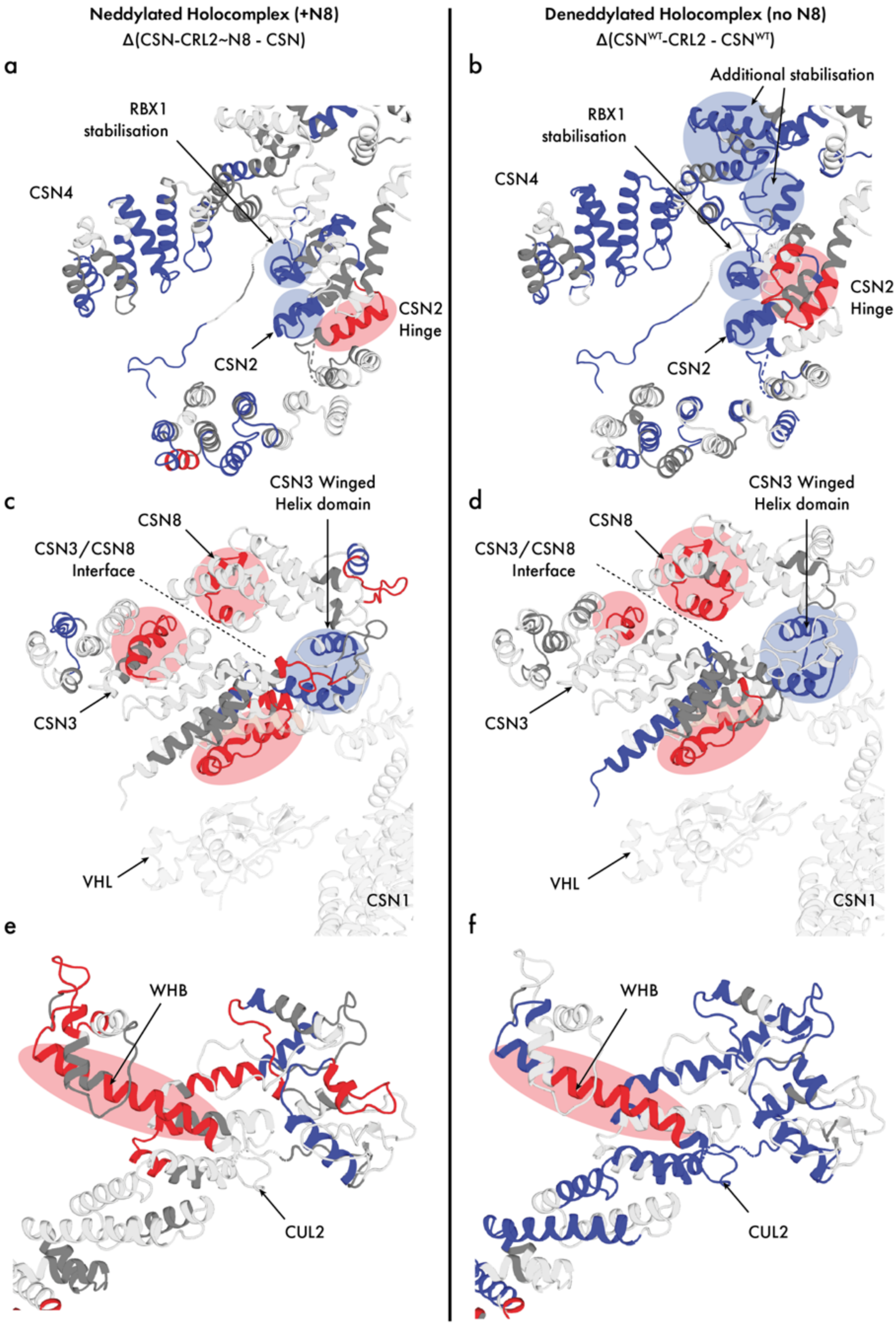

Neddylated Holocomplex (+N8)
Δ(CSN-CRL2~N8 - CSN)
Deneddylated Holocomplex (no N8)
Δ(CSN^WT-CRL2 - CSN^WT)
a
RBX1
stabilisation
CSN4
CSN2
Hinge
CSN2
b
Additional stabilisation
RBX1
stabilisation
CSN4
CSN2
Hinge
CSN2
c
CSN3 Winged
Helix domain
CSN8
CSN3/CSN8
Interface
CSN3
VHL
CSN1
d
CSN3 Winged
Helix domain
CSN8
CSN3/CSN8
Interface
CSN3
VHL
CSN1
e
WHB
CUL2
f
WHB
CUL2

**Fig. S12. ΔHDX changes in CSN2/CSN4/RBX1, CSN3/CSN8 and CUL2.** Regions showing significant destabilisation (red) and stabilisation (blue) are shown on the structures of CSN-CRL2~N8 (left) and CSN-CRL2 (right) and highlighted for clarity. (**a-b**) Stabilisation of CSN2 and RBX1 interfaces. In the deneddylated complex, CSN4 and RBX1 stabilisation suggests an interface between the two subunits (**b**; top right blue circles). (**c-d**) HDX changes in CSN3, CSN8. VHL and CSN1 (shown in white) are displayed for reference. Both CSN3 and CSN8 experience destabilisation at their interfaces in both left and right complexes. The CSN3 surface closest to VHL (red) and the CSN3 winged helix domain (blue) also shows consistent differences. (**e-f**) Destabilisation of the WHB domain Cullin-2 (CUL2) in both conditions.

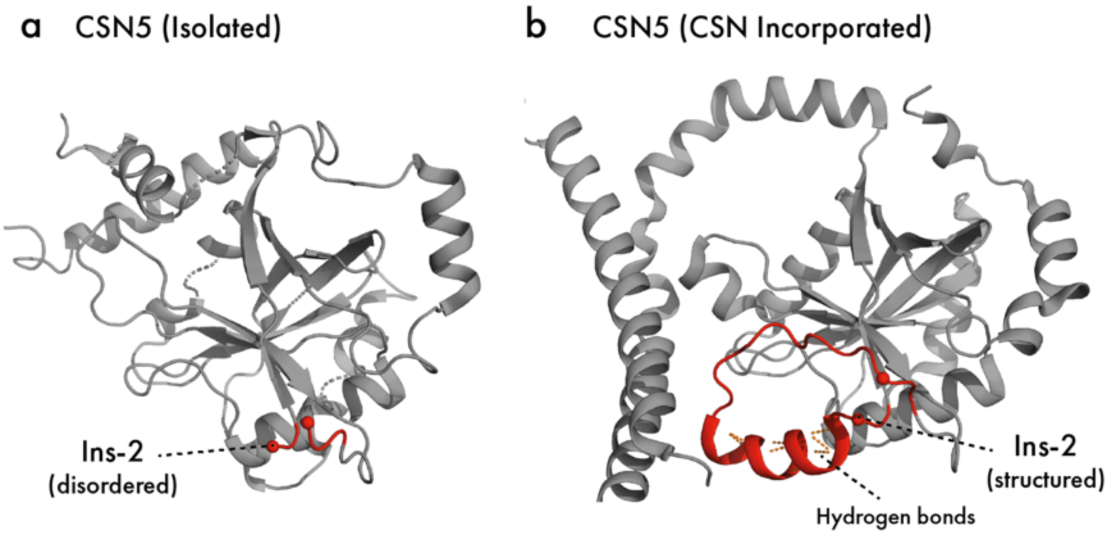


**Fig. S13. CSN5 Ins-2 loop in isolated and CSN incorporated structures.** (**a**) Disordered Ins-2 loop in isolated CSN (PDB 4F7O). (**b**) Structured Ins-2 loop in CSN incorporated CSN5 (PDB 4D10). Hydrogen bonds of the Ins-2 helix have been shown in orange.

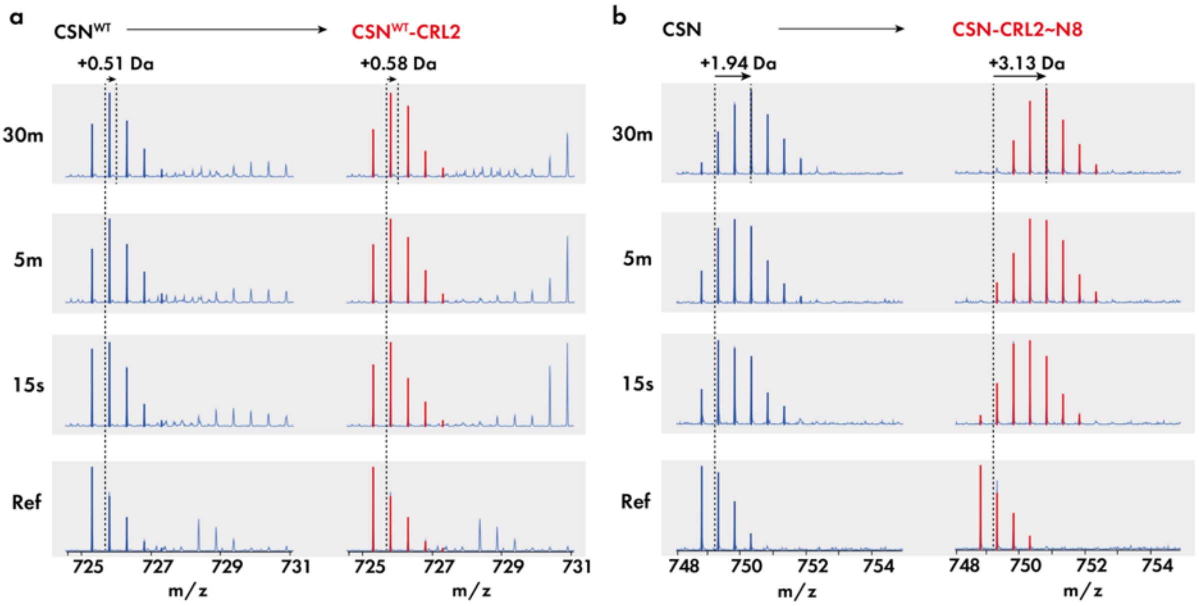


**Fig. S14. Deuteration profiles of CSN5 active site.** Profiles of the (**a**) non-neddylated and (**b**) neddylated holocomplex (red) are compared with deuteration profiles of the isolated CSN (blue). The ion distribution of active site peptide $^{134}$GWYXSHPGYGCW$^{145}$ (where X indicates H in CSN$^{WT}$ and A in CSN$^{5H138A}$) is shown for (a) and (b) across reference (0s), 15s, 5m and 30m timepoints. The mass of the non-deuterated reference peptide is shown by the dotted line. A second dotted line at time 30m for the holocomplexes (red) indicates the mass of the peptide at time 30m. The relative deuterium change of the active site peptide in (a) following binding of CRL2 is negligible, while in (b), presence of CRL2~N8 leads to an increase of mass.

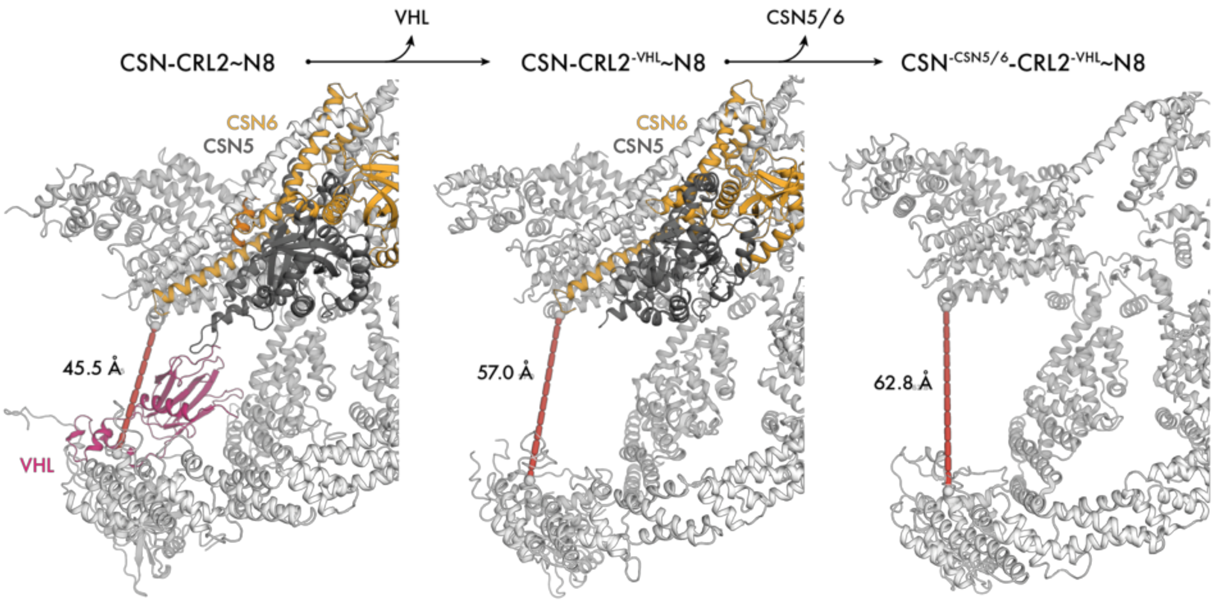


**Fig. S15. Conformational heterogeneity of the CSN-CRL2~N8 structures.** The distance between CSN3 and CUL2 N-terminal domain of CSN-CRL2~N8, after loss of VHL and after loss of CSN5/CSN6 is shown by the dashed red line. All other CSN-CRL2 subunits have been coloured in white for clarity.

### Supplementary Tables

##### Table S1. Chemical cross-links of CSN-CRL2~N8

| Index | Protein A | Protein B | Residue A | Residue B |
|---|---|---|---|---|
| | | **Intra-subunit** | | |
| 1 | CSN1 | CSN1 | 183 | 153 |
| 2 | CSN1 | CSN1 | 444 | 451 |
| 3 | CSN1 | CSN1 | 144 | 136 |
| 4 | CSN1 | CSN1 | 343 | 332 |
| 5 | CSN2 | CSN2 | 150 | 157 |
| 6 | CSN2 | CSN2 | 110 | 150 |
| 7 | CSN2 | CSN2 | 253 | 215 |
| 8 | CSN2 | CSN2 | 415 | 364 |
| 9 | CSN2 | CSN2 | 426 | 364 |
| 10 † | CSN1 | CSN1 | 467 | 467 |
| 11 | CSN2 | CSN2 | 243 | 157 |
| 12 | CSN2 | CSN2 | 56 | 43 |
| 13 | CSN2 | CSN2 | 64 | 93 |
| 14 | CSN2 | CSN2 | 81 | 48 |
| 15 | CSN2 | CSN2 | 48 | 64 |
| 16 | CSN2 | CSN2 | 48 | 93 |
| 17 | CSN2 | CSN2 | 93 | 73 |
| 18 | CSN2 | CSN2 | 64 | 73 |
| 19 † | CSN2 | CSN2 | 415 | 415 |
| 20 | CSN3 | CSN3 | 254 | 243 |
| 21 | CSN3 | CSN3 | 125 | 115 |
| 22 | CSN3 | CSN3 | 254 | 281 |
| 23 | CSN4 | CSN4 | 227 | 214 |
| 24 | CSN4 | CSN4 | 214 | 188 |
| 25 | CSN5 | CSN5 | 308 | 179 |
| 26 | CSN5 | CSN5 | 193 | 218 |
| 27 | CSN5 | CSN5 | 218 | 294 |
| 28 | CSN5 | CSN5 | 218 | 298 |
| 29 | CSN6 | CSN6 | 75 | 113 |
| 30 | CSN6 | CSN6 | 229 | 236 |
| 31* | CSN7b | CSN7b | 221 | 217 |
| 32* | CSN7b | CSN7b | 221 | 257 |
| 33* | CSN7b | CSN7b | 217 | 199 |
| 34* | CSN7b | CSN7b | 218 | 217 |
| 35 | CUL2 | CUL2 | 393 | 382 |
| 36 | CUL2 | CUL2 | 179 | 252 |
| 37 | CUL2 | CUL2 | 393 | 404 |
| 38 | CUL2 | CUL2 | 677 | 720 |

| | | | | |
|---|---|---|---|---|
| 39 | CUL2 | CUL2 | 720 | 382 |
| 40 | CUL2 | CUL2 | 728 | 677 |
| 41 | CUL2 | CUL2 | 252 | 244 |
| 42 | CUL2 | CUL2 | 433 | 386 |
| 43 | CUL2 | CUL2 | 608 | 615 |
| 44 | CUL2 | CUL2 | 238 | 244 |
| 45 | CUL2 | CUL2 | 404 | 400 |
| 46 | CUL2 | CUL2 | 433 | 677 |
| 47 | CUL2 | CUL2 | 62 | 149 |
| 48 | CUL2 | CUL2 | 179 | 96 |
| 49 | CUL2 | CUL2 | 433 | 472 |
| 50 | CUL2 | CUL2 | 677 | 382 |
| 51 | ELOB | ELOB | 19 | 36 |
| 52 | ELOB | ELOB | 36 | 28 |
| 53 | ELOB | ELOB | 36 | 46 |
| 54 | ELOB | ELOB | 11 | 28 |
| 55 | ELOB | ELOB | 11 | 36 |
| 56 | ELOB | ELOB | 46 | 28 |
| 57 † | ELOB | ELOB | 104 | 104 |
| 58 | NEDD8 | NEDD8 | 60 | 6 |
| 59 | NEDD8 | NEDD8 | 11 | 33 |
| 60 | VHL | VHL | 196 | 171 |
| | | **Inter-subunit** | | |
| 1 | CSN1 | CSN2 | 438 | 426 |
| 2 | CSN1 | CSN2 | 438 | 415 |
| 3* | CSN1 | CSN3 | 467 | 406 |
| 4 | CSN2 | CSN3 | 426 | 243 |
| 5 | CSN2 | CSN4 | 361 | 337 |
| 6 | CSN2 | CSN4 | 376 | 337 |
| 7 | CSN2 | CSN7b | 426 | 199 |
| 8 | CSN3 | CSN8 | 296 | 58 |
| 9 | CSN3 | CSN8 | 115 | 65 |
| 10 | CSN3 | CSN8 | 115 | 58 |
| 11 | CSN5 | CSN7b | 308 | 199 |
| 12 | CSN5 | CSN7b | 179 | 199 |
| 13* | CSN5 | CSN7b | 294 | 217 |
| 14 | CUL2 | NEDD8 | 382 | 33 |
| 15 | CUL2 | NEDD8 | 433 | 6 |
| 16 | CUL2 | VHL | 114 | 196 |
| 17 | CUL2 | ELOC | 96 | 43 |
| 18 | CUL2 | ELOC | 114 | 43 |
| | | **Inter-complex** | | |
| 19 | CSN1 | ELOB | 158 | 19 |
| 20 | CSN2 | CUL2 | 263 | 462 |

| | | | | |
|---|---|---|---|---|
| 21 | CSN2 | CUL2 | 157 | 489 |
| 22 | CSN2 | CUL2 | 225 | 462 |
| 23 | CSN2 | CUL2 | 64 | 404 |
| 24 | CSN4 | RBX1 | 200 | 105 |

* indicates six cross-links omitted from projection onto the CSN-CRL2~N8 structure in **Fig. S5b-c** due to lack of structural coverage.

† denotes three intra-subunit cross-links to the same residue. Also omitted from **Fig. S5b-c**.

**Table S2. Chemical cross-links of CSN-CRL2**

| Index | Protein A | Protein B | Residue A | Residue B |
|---|---|---|---|---|
| | | **Intra-subunit** | | |
| 1 | CSN1 | 183 | CSN1 | 153 |
| 2 | CSN1 | 444 | CSN1 | 451 |
| 3 | CSN2 | 64 | CSN2 | 93 |
| 4 | CSN2 | 110 | CSN2 | 150 |
| 5 | CSN2 | 225 | CSN2 | 189 |
| 6 | CSN2 | 225 | CSN2 | 253 |
| 7 | CSN2 | 253 | CSN2 | 215 |
| 8 | CSN2 | 253 | CSN2 | 300 |
| 9 | CSN2 | 415 | CSN2 | 364 |
| 10 | CSN2 | 426 | CSN2 | 415 |
| 11 | CSN2 | 150 | CSN2 | 157 |
| 12 | CSN2 | 157 | CSN2 | 243 |
| 13 | CSN2 | 48 | CSN2 | 81 |
| 14 | CSN2 | 48 | CSN2 | 93 |
| 15 | CSN3 | 125 | CSN3 | 115 |
| 16 | CSN3 | 254 | CSN3 | 243 |
| 17 | CSN3 | 254 | CSN3 | 281 |
| 18 | CSN4 | 214 | CSN4 | 200 |
| 19 | CSN4 | 150 | CSN4 | 200 |
| 20 | CSN4 | 188 | CSN4 | 214 |
| 21 | CSN5 | 191 | CSN5 | 219 |
| 22 | CSN5 | 194 | CSN5 | 219 |
| 23 | CSN5 | 180 | CSN5 | 309 |
| 24* | CSN7b | 221 | CSN7b | 217 |
| 25 | CUL2 | 62 | CUL2 | 149 |
| 26 | CUL2 | 179 | CUL2 | 96 |
| 27 | CUL2 | 179 | CUL2 | 252 |
| 28 | CUL2 | 252 | CUL2 | 244 |
| 29 | CUL2 | 261 | CUL2 | 244 |
| 30 | CUL2 | 393 | CUL2 | 404 |
| 31 | CUL2 | 608 | CUL2 | 615 |
| 32 | CUL2 | 659 | CUL2 | 404 |
| 33 | CUL2 | 244 | CUL2 | 238 |
| 34 | CUL2 | 400 | CUL2 | 404 |
| 35 | CUL2 | 22 | CUL2 | 74 |
| 36 | CUL2 | 290 | CUL2 | 291 |
| 37 | CUL2 | 244 | CUL2 | 291 |
| 38 | CUL2 | 400 | CUL2 | 401 |
| 39 | CUL2 | 677 | CUL2 | 728 |

| | | | | |
|---|---|---|---|---|
| 40 | CUL2 | 719 | CUL2 | 728 |
| 41[†] | CUL2 | 252 | CUL2 | 252 |
| 42 | ELOB | 36 | ELOB | 46 |
| 43 | ELOB | 11 | ELOB | 28 |
| 44 | ELOB | 11 | ELOB | 36 |
| 45 | ELOB | 11 | ELOB | 46 |
| 46 | ELOB | 28 | ELOB | 36 |
| 47 | VHL | 196 | VHL | 171 |
| | | **Inter-subunit** | | |
| 1 | CSN1 | 422 | CSN2 | 415 |
| 2 | CSN1 | 438 | CSN2 | 426 |
| 3 | CSN1 | 438 | CSN3 | 312 |
| 4* | CSN1 | 467 | CSN7b | 257 |
| 5 | CSN2 | 303 | CSN4 | 200 |
| 6 | CSN2 | 426 | CSN3 | 243 |
| 7 | CSN2 | 361 | CSN4 | 337 |
| 8 | CSN3 | 115 | CSN8 | 58 |
| 9 | CSN3 | 115 | CSN8 | 65 |
| 10 | CSN4 | 372 | CSN6 | 75 |
| 11 | CSN5 | 180 | CSN7b | 199 |
| 12 | CSN5 | 309 | CSN7b | 199 |
| 13* | CSN5 | 294 | CSN7b | 217 |
| 14 | CUL2 | 114 | VHL | 196 |
| 15 | CUL2 | 114 | ELOC | 43 |
| 16 | ELOB | 19 | ELOC | 32 |
| 17 | ELOB | 36 | ELOC | 20 |
| 18 | ELOB | 19 | ELOC | 6 |
| | | **Inter-complex** | | |
| 19 | CSN1 | 467 | VHL | 171 |
| 20 | CSN2 | 225 | CUL2 | 462 |
| 21 | CSN2 | 263 | CUL2 | 462 |
| 22 | CSN2 | 415 | CUL2 | 692 |
| 23* | CSN7b | 218 | CUL2 | 149 |
| 24* | CSN7b | 221 | CUL2 | 291 |

* indicates five cross-links omitted from projection onto the CSN-CRL2 structure in **Fig. S8b-c** due to lack of structural coverage.

[†] denotes one intra-subunit cross-links to the same residue. Also omitted from **Fig. S8b-c**.

#### Supplementary References


1 Scheres, S. H. RELION: implementation of a Bayesian approach to cryo-EM structure determination. *J Struct Biol* **180**, 519-530, doi:10.1016/j.jsb.2012.09.006 (2012).
